# Highly repeatable phenotypic consequences of whole-genome duplication in *Spirodela polyrhiza*

**DOI:** 10.64898/2026.01.23.701260

**Authors:** Mortier Frederik, Quinten Bafort, Bonte Dries, Yves Van de Peer

## Abstract

Whole-genome duplication (WGD) is widespread in plants, yet the extent to which it yields predictable phenotypic outcomes remains unclear. Here we show that the phenotypic consequences of genome doubling in a duckweed model system, *Spirodela polyrhiza*, are highly repeatable and largely deterministic. We previously generated three independent colchicine-induced autotetraploids from each of nine globally distributed diploid genotypes and, here, quantified growth and morphology across a salt gradient. In benign conditions, diploids grew faster, whereas tetraploids had larger, thicker fronds; with increasing salinity, the diploid advantage diminished and tetraploids often matched or exceeded diploid growth. Variance partitioning revealed that ploidy *per se* explains more phenotypic variation, including salt tolerance, than genotypic background, with rare within-genotype stochasticity. These results indicate that the shifts in morphology and stress tolerance from genome doubling are predictable and can outweigh the phenotypic effect from genetic sequence diversity.

## Introduction

Polyploidy, the result of Whole Genome Duplication (WGD), has repeatedly shaped plant diversification and ecological success (Soltis *et al*., 2009). WGD can frequently create “hopeful monsters”, the product of a large phenotypic change that “hopes” for an evolutionary context in which it is not selected against (Otto and Whitton, 2000; Dietrich, 2003; Ebadi *et al*., 2023). Evidence for the phenotypic effects of polyploidy is largely based on patterns across taxa and environments (e.g., Van De Peer, Mizrachi and Marchal, 2017; Bomblies, 2020; Clo and Kolář, 2021; Van de Peer *et al*., 2021; Mortier *et al*., 2024), leaving open how much of the polyploid phenotype is predictable versus stochastic. Addressing this requires isolating WGD per se in autopolyploids from other genetic mechanisms, such as genotypic background and hybridization, and experimentally quantifying its contribution to phenotypic heterogeneity across environments.

Polyploidy can affect a wide range of traits. From a quantitative genetics perspective, phenotypic heterogeneity is caused by genetic (i.e., heritable) factors and environmental influences, to which polyploidy adds an extra layer of genetic complexity. The only trait consistently associated with genome doubling across all polyploid plant species is an increase in cell size (Ramsey and Schemske, 2002; Bomblies, 2020). Other reported effects include slower growth rates but larger overall plant and organ size, a higher prevalence of asexual reproduction or selfing, and fewer but larger stomata. Furthermore, WGD can trigger transcriptomic and epigenetic remodelling (e.g., as dosage compensation mechanism) and changes in nuclear architecture (Doyle and Coate, 2019; Song *et al*., 2020; Birchler and Veitia, 2021), all of which are heritable with respect to the polyploid state even when not driven by coding sequence changes. Polyploidy’s association with an enhanced tolerance to stressful environmental conditions, furthermore, affects how the genetic component of phenotypic heterogeneity interacts with the environmental influence (GxE interactions). For each of these associations with polyploidy, however, numerous counterexamples have been published (many in Bomblies, 2020).

Within species, induced polyploids often show genotype-specific responses (Husband, Baldwin and Sabara, 2016; Anneberg and Segraves, 2020; Ponsford *et al*., 2022; Anneberg *et al*., 2023; Mattingly and Hovick, 2023), and, even within a single genotype, independent polyploid lines can vary in cellular and physiological traits (Yu *et al*., 2010; Aversano *et al*., 2013, but see Dixit, Verma and Chaudhary, 2015; Zhang *et al*., 2015; Zhou *et al*., 2015 and Chen *et al*., 2020 where they do not). However, key quantitative traits such as macromorphological traits, growth, and other fitness measurements tend to vary considerably more between independent polyploid lines of the same progenitor genotype (Kulkarni and Borse, 2010; Aversano *et al*., 2013; Münzbergová, 2017; Tan *et al*., 2017; Corneillie *et al*., 2019; Wei *et al*., 2020, but see Dixit, Verma and Chaudhary, 2015; Chen *et al*., 2020; Jiang *et al*., 2020). These observations suggest that the polyploid phenotype comprises both a deterministic component (repeatable consequences of genome doubling) and a stochastic component, whose magnitudes remain largely unquantified.

To quantify these components, replication is required at three levels: (i) multiple genotypic backgrounds, (ii) independent genome duplication events within each genotype, and (iii) replicated measurements per strain (Figure 1). If WGD-induced phenotypic changes are deterministic, their consequence will be repeatable across independent polyploids, either across genotypes or in a genotype-specific manner. Conversely, if WGD introduces stochastic effects, the average phenotypic outcome will vary unpredictably among independently derived polyploids from the same progenitor. Compared to most studies who lack at least one level of replication to distinguish all sources of possible variation (to our knowledge, only Aversano *et al*., 2013 show the required replication), our clonal model permitted true biological replication of independent autotetraploids in fixed genotypic backgrounds, avoiding confounding from meiotic chromosome reshuffling. We therefore generated three independent autotetraploids from each of nine diploid genotypes spanning global diversity in Spirodela polyrhiza and measured growth and morphology across a salt gradient. This design enabled variance partitioning into effects of ploidy, genotype, their interaction, line-specific (stochastic) variation, and residual sources (Figure 1). We measured growth rate in number, biomass, and surface area of these 36 strains across a salt gradient with 5 replicated measurements each. Tetraploidy was artificially induced using an antimitotic agent, colchicine, ensuring that both subgenomes were inherited from the parent diploid to avoid effects of subgenome differentiation.

**Figure 1.**
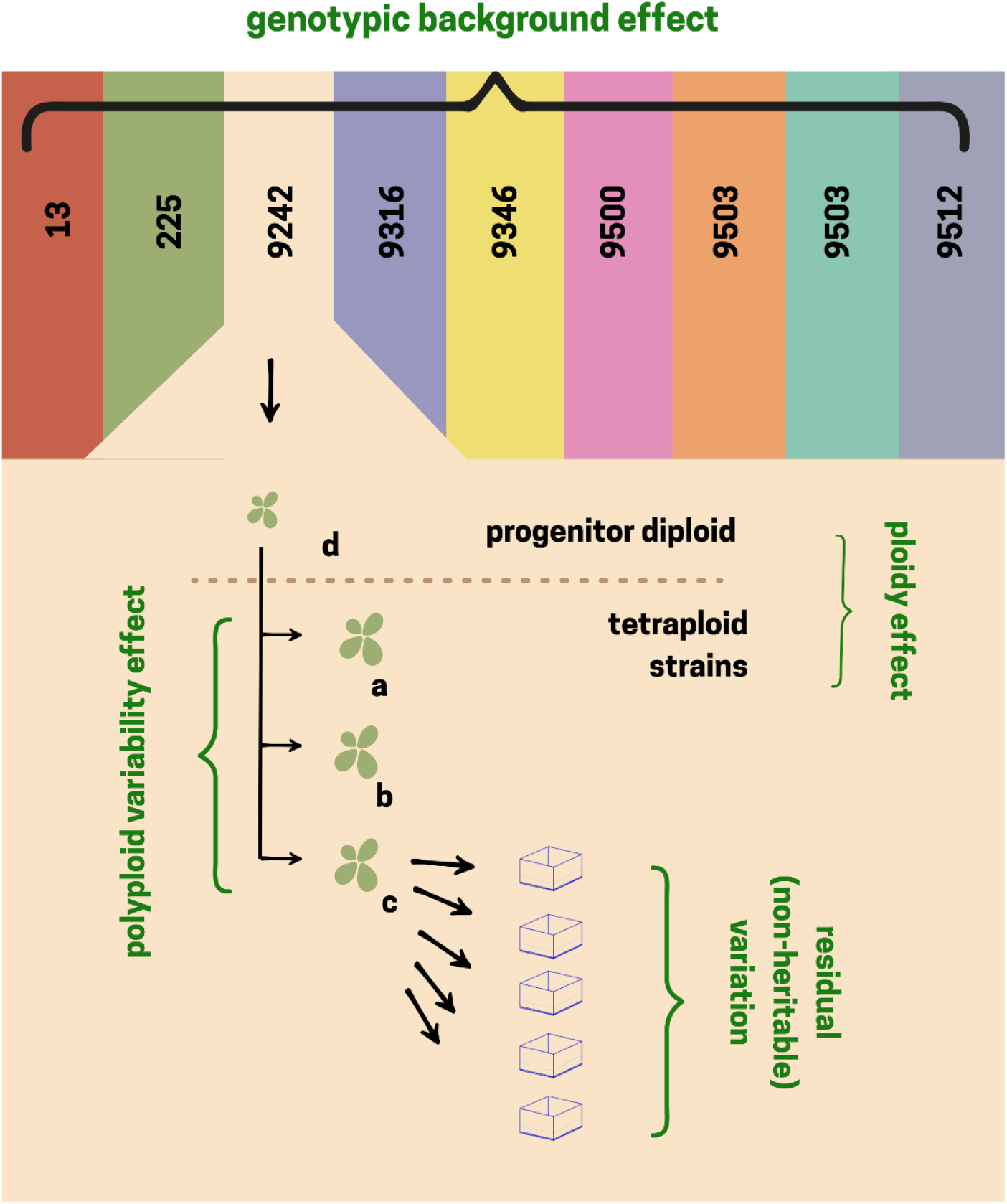
Heterogeneity in growth rate and morphology in artificial tetraploid Spirodela polyrhiza populations is affected by the deterministic component of the ploidy effect, but also a stochastic component of the ploidy effect (polyploid variability), the genotype, the genotype-specific ploidy effect (interaction genotype X ploidy) and residual variability that stems from non-heritable phenotypic heterogeneity, such as regulatory noisiness or auxiliary environmental fluctuations, and measurement error. To disentangle all these sources, we partitioned the variance in measured outcomes from multiple genotypic backgrounds (codes and colors on top for each of nine backgrounds) with each multiple artificial tetraploid strains (a, b and c) compared with their diploid progenitor (d) and five replicated measurements per strain (represented by five growing containers).

## Results

### Growth in control conditions

We observed a reduced growth rate in the absence of salt stress (control) in 25 out of 27 tetraploids compared to their progenitor diploid strain (Figure 2), although growth rates varied substantially across genotypes. We show growth in fresh weight here (Figure 2) and in further results but observed the same trend in growth in frond count and surface area (Figure S4). Tetraploid strain 9512b, one of the two strains that did not show a reduced growth rate, was later found to be mixoploid (Figure S3) and therefore removed from further analysis.

**Figure 2.**
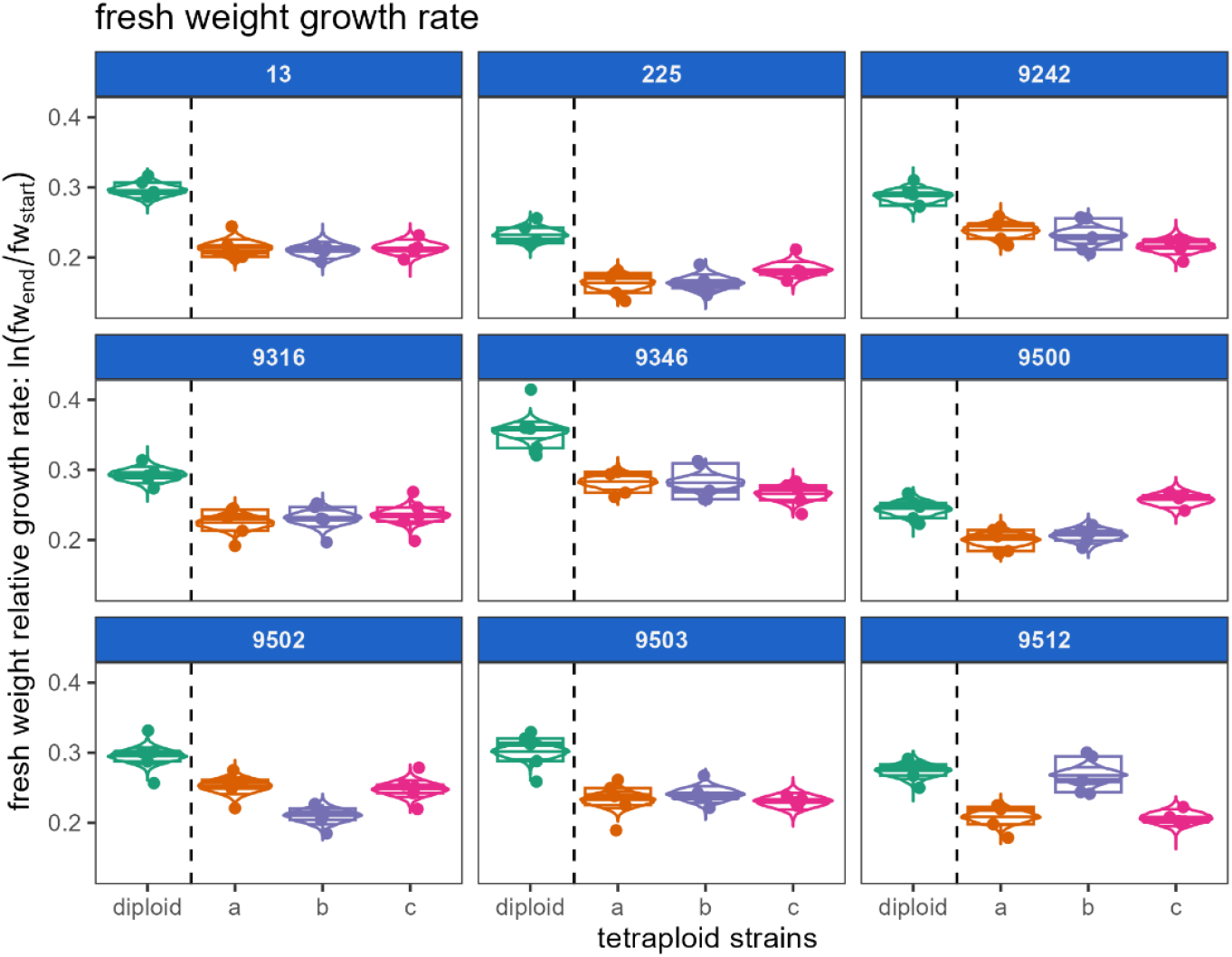
relative growth rate (RGR) in control conditions in terms of fresh of three independent colchitetraploid strains (a, b, c) compared to their progenitor diploid strain for nine different genotypic backgrounds. Dots and boxplots show the observations and violins indicate the posterior expected RGR for that strain with the 9th, 50th and 91st percentile indicated. We show the results for the mixoploid strain 9512b but exclude it for subsequent analyses.

When partitioning variability in fresh weight, we found that the genotype-specific variability among replicate tetraploid strains (representing the stochastic component) was generally lower than the deterministic effect of ploidy, strain, and their interaction (Figure 3). We found the same low amount of variability in growth in frond count and surface area that is explained by replicated tetraploids (Figure S5). Notable exceptions occur in genotypes 9500 and 9502, driven by lines 9500c, which showed diploid-like growth, and 9502b with exceptionally slow growth.

**Figure 3.**
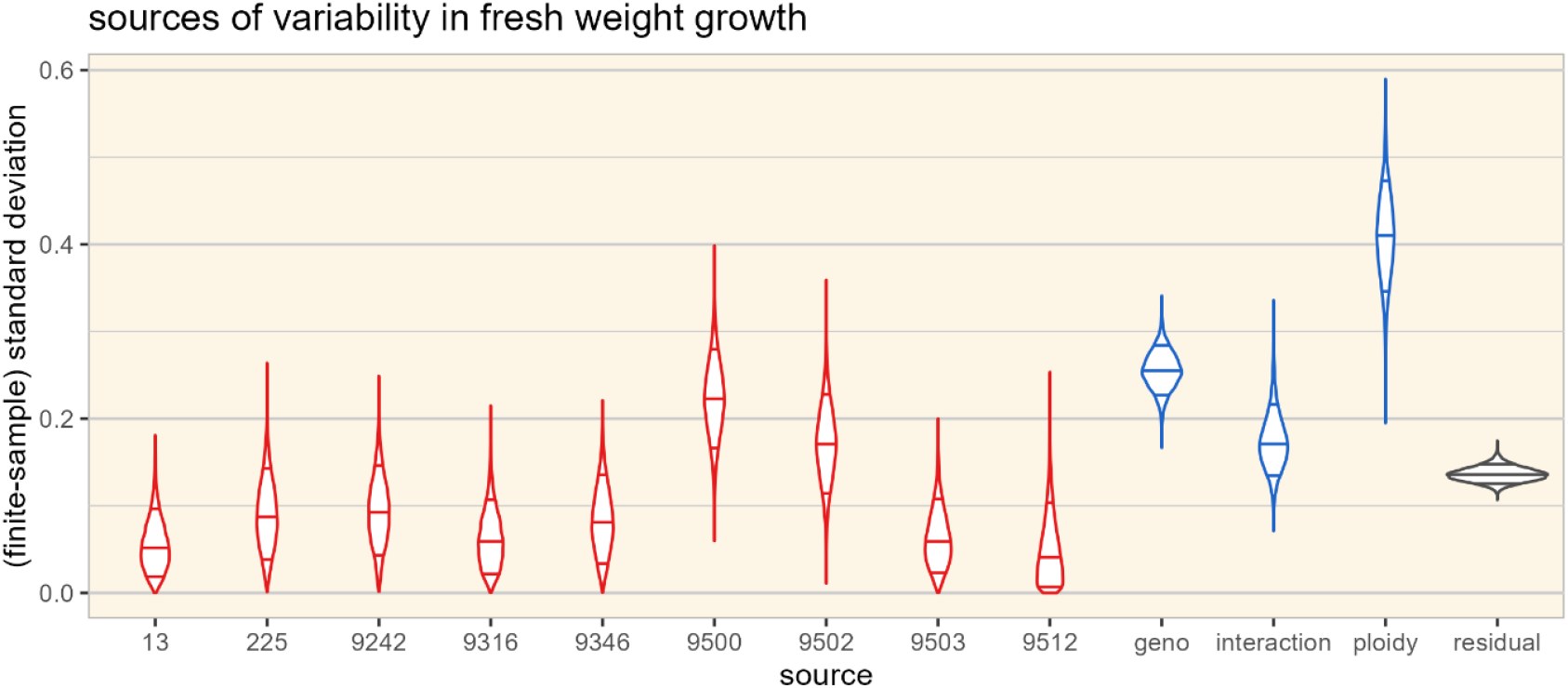
Posterior distribution of finite-sample standard deviation of all coefficients within each effect for the model of growth of fresh weight in control conditions. This shows the variation estimated by the model caused by stochastic polyploidy effects that differ between independent polyploidy events (strain-specific, in red) compared to the deterministic effects of ploidy, genotype (geno), their interaction and the remaining variation among repeated measurements (residual, black). We indicated the 9th, 50th and 91th percentile in each violin.

Growth in number of fronds, fresh weight and frond surface area showed strong correlations (Figure S6). This conservative relation reflects a consistent fresh weight per individual, average frond surface area and specific leaf area (frond leaf surface area, here per unit of fresh weight) within each genotype. We observed elevated individual fresh weight and frond surface area alongside decreased specific leaf area in all tetraploid strains except strain 9500c compared to their respective progenitor diploid (Figure 4). However, correlations with growth rate were only evident between strains, not within replicate measurements of the same strain (Figure S7). This suggests a genotypic basis for the observed variation rather than a biophysical constraint from polyploidy.

**Figure 4.**
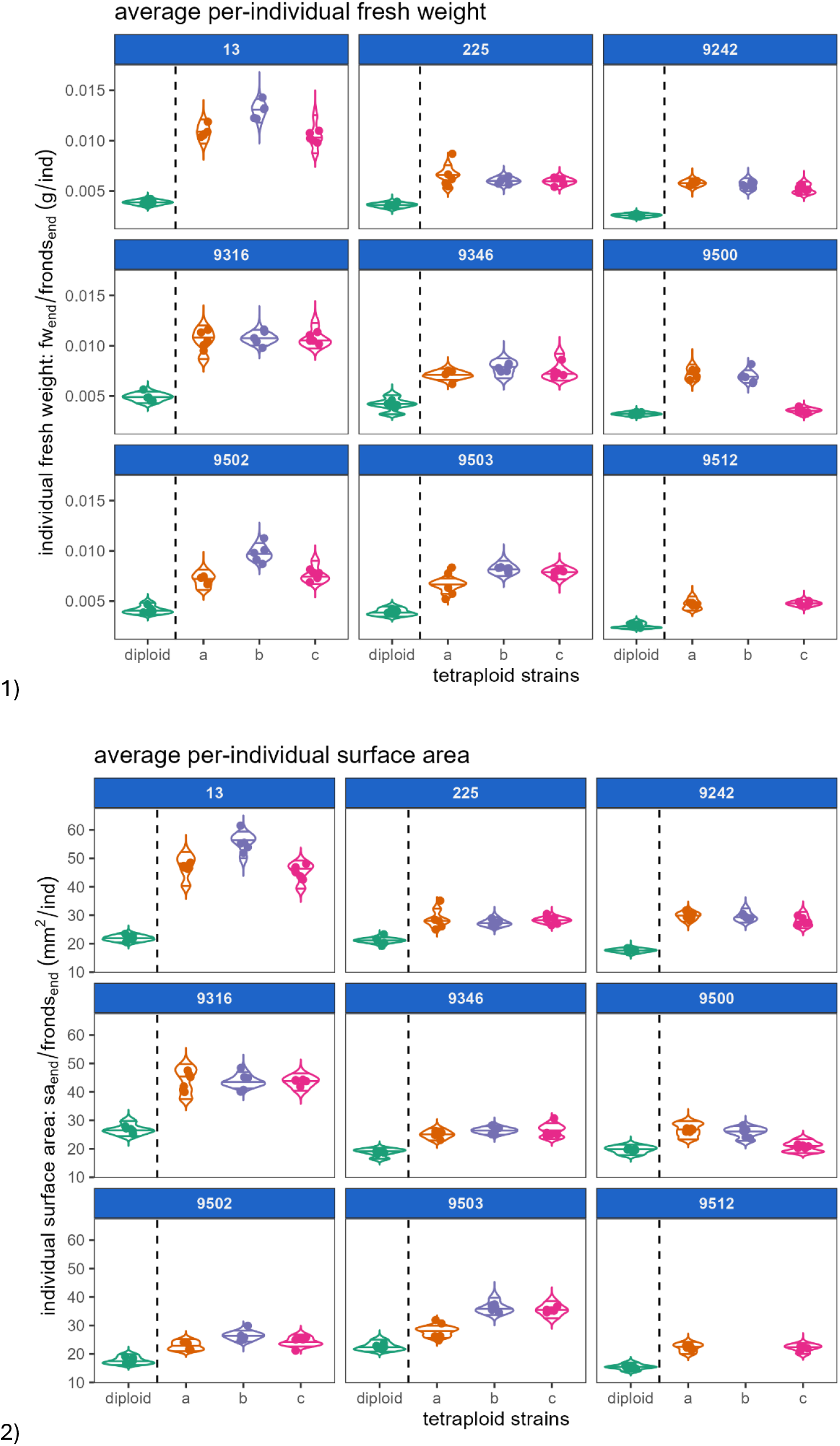
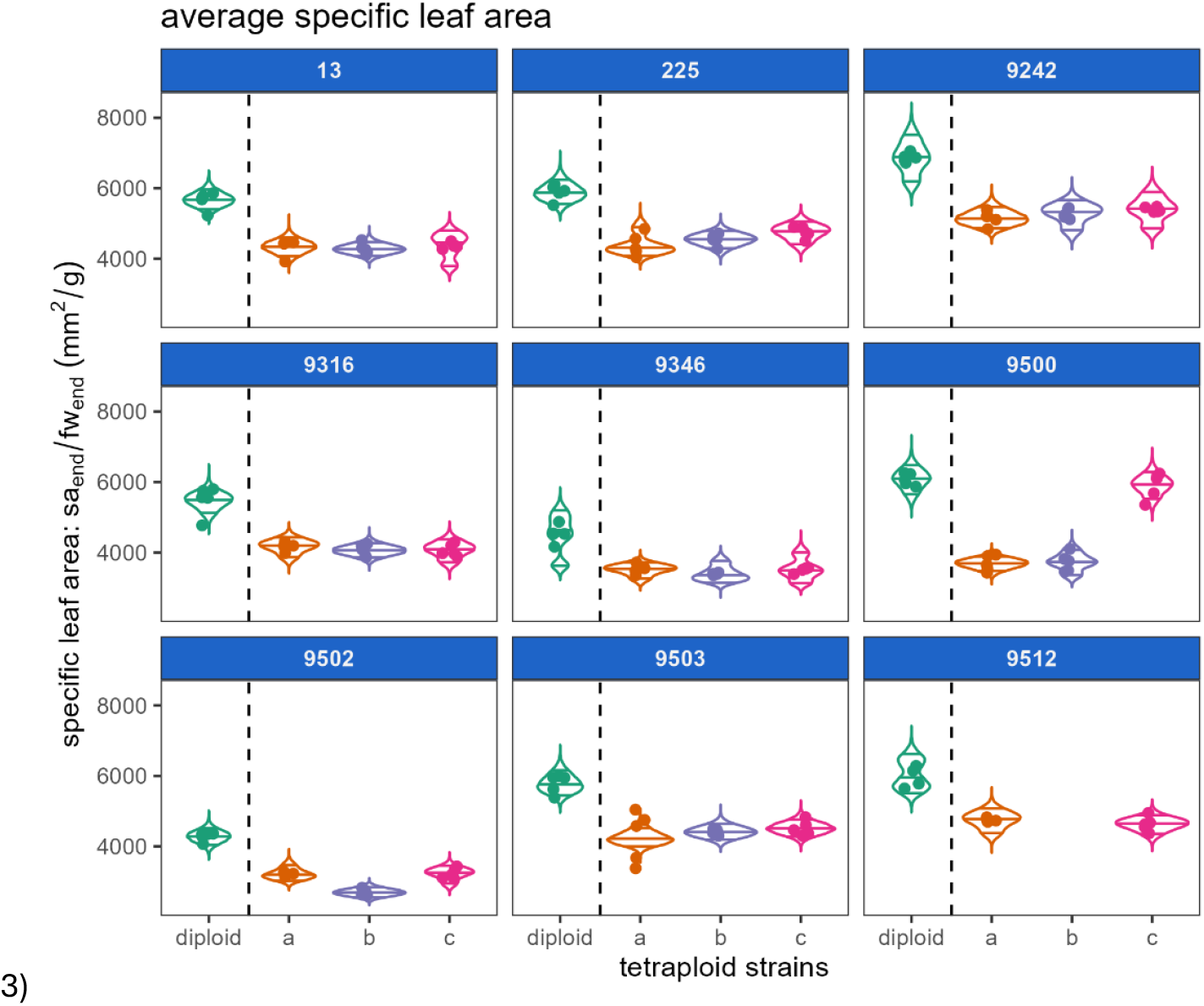
average morphological traits in control conditions: 1) average individual fresh weight, 2) average frond surface area and 3) average fresh weight specific leaf area of three independent colchitetraploid strains (a, b, c) compared to their progenitor diploid strain for nine different genotypic backgrounds. Dots and boxplots show the observations and violins indicate the posterior expected RGR for that strain with the 9th, 50th and 91st percentile indicated.

### Growth in salt gradient

Fresh weight growth of all strains, across all genotypes, ploidy levels, and growth metrics decreased with an increasing salt concentration (Figure 5). Notably, the diploid advantage over its descendant tetraploid strains diminished as salt concentration increased. In some cases, tetraploid strains even outperformed their diploid progenitor at higher salt concentrations (Figure 5, Figure S8-9). Growth in frond count and surface area similarly decreased with salt and similarly decreased slower in tetraploid compared to their progenitor diploid strain (Figure S8-10). For example, tetraploid 9242b showed superior performance across all three metrics at 8g/l NaCl.

**Figure 5.**
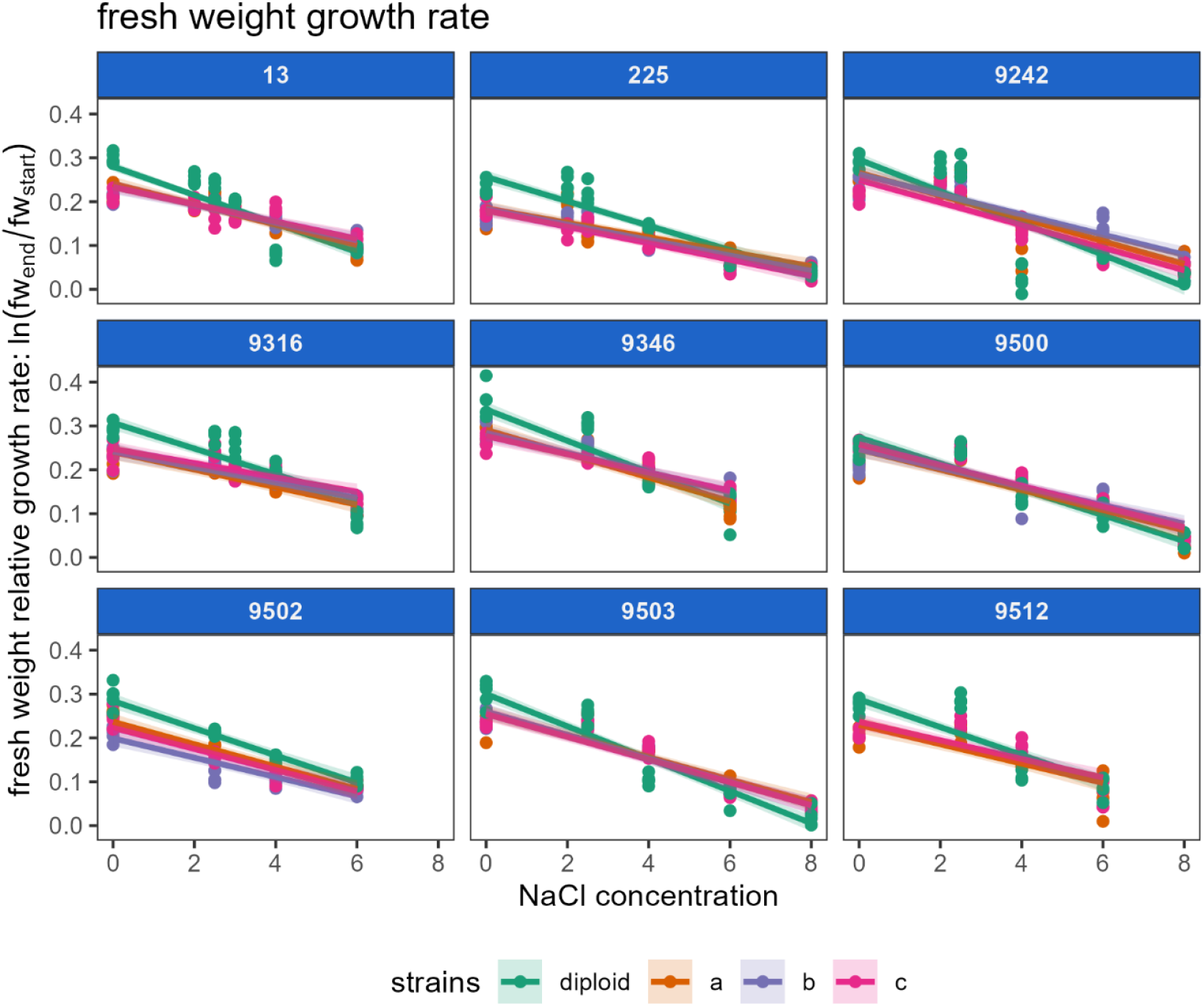
Relative growth rate in fresh weight of three independent colchitetraploid strains (a, b, c) compared to their progenitor diploid strain (in green) for nine different genotypic backgrounds. Dots show the observations and lines indicate the median of the posterior expected RGR for that strain with the 4th and 96th percentile indicated by the ribbon.

Variability in growth across the salt gradient is foremostly affected by the salt concentration, independent of ploidy and genotype (Figure 6). This reaction norm (slope that represents the salt response) is additionally affected by ploidy, corresponding with the consistent slower decrease in tetraploid growth rates with increasing salt concentration compared to diploids. Just as in the control tests only, replicated tetraploid duckweed strains varied only to a limited extent within each genotype compared to the other sources. This lack of variability between replicated tetraploids held true for both the intercept (*α*) and slope across the salt gradient (i.e., reaction norm *β*) for variability and also for growth in frond count (Figure S11). For growth in surface area, we estimated a weaker deterministic effect of ploidy on salt tolerance, comparable to the other sources. In a few genotypes, the model estimates elevated variability among repeated polyploidy events within each genotype that were inconsistent across growth metrics.

**Figure 6.**
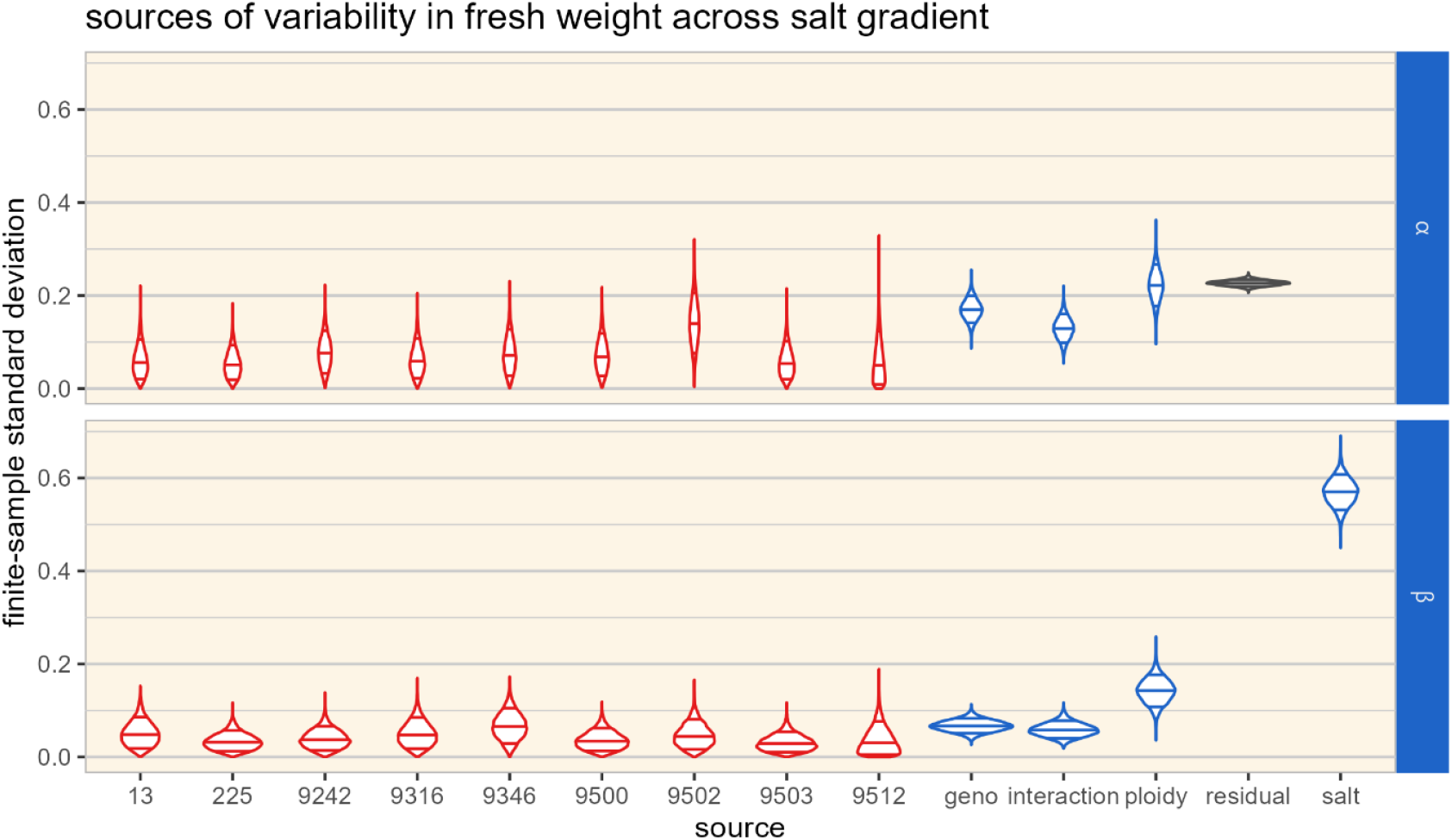
Upper panel α: estimated variability components affecting the intercept for growth in fresh weight. Lower panel β: estimated variability components affecting the salt stress response of fresh weight growth. Both panels show the finite-sample standard variation of all statistical model coefficients of the stochastic strain-specific WGD effects (red) compared to the deterministic ploidy effect, genotype (geno), their interaction and the residual variance (residual, black). The “salt” component shows the variation in outcome caused by the overall slope across salt concentration on the diploid, independent of any genotype or strain effect. We indicated the 9th, 50th and 91st percentile in each violin.

We observed that individual traits were affected by increasing salt concentrations. Individual fresh weight, frond surface area and specific leaf area decreases with increasing salt concentration (Figure 7). More detailed analyses on how the correlation between individual traits and growth rate shifts across the salt gradient are provided in supplementary materials (Appendix 12, Figure S12)

**Figure 7.**
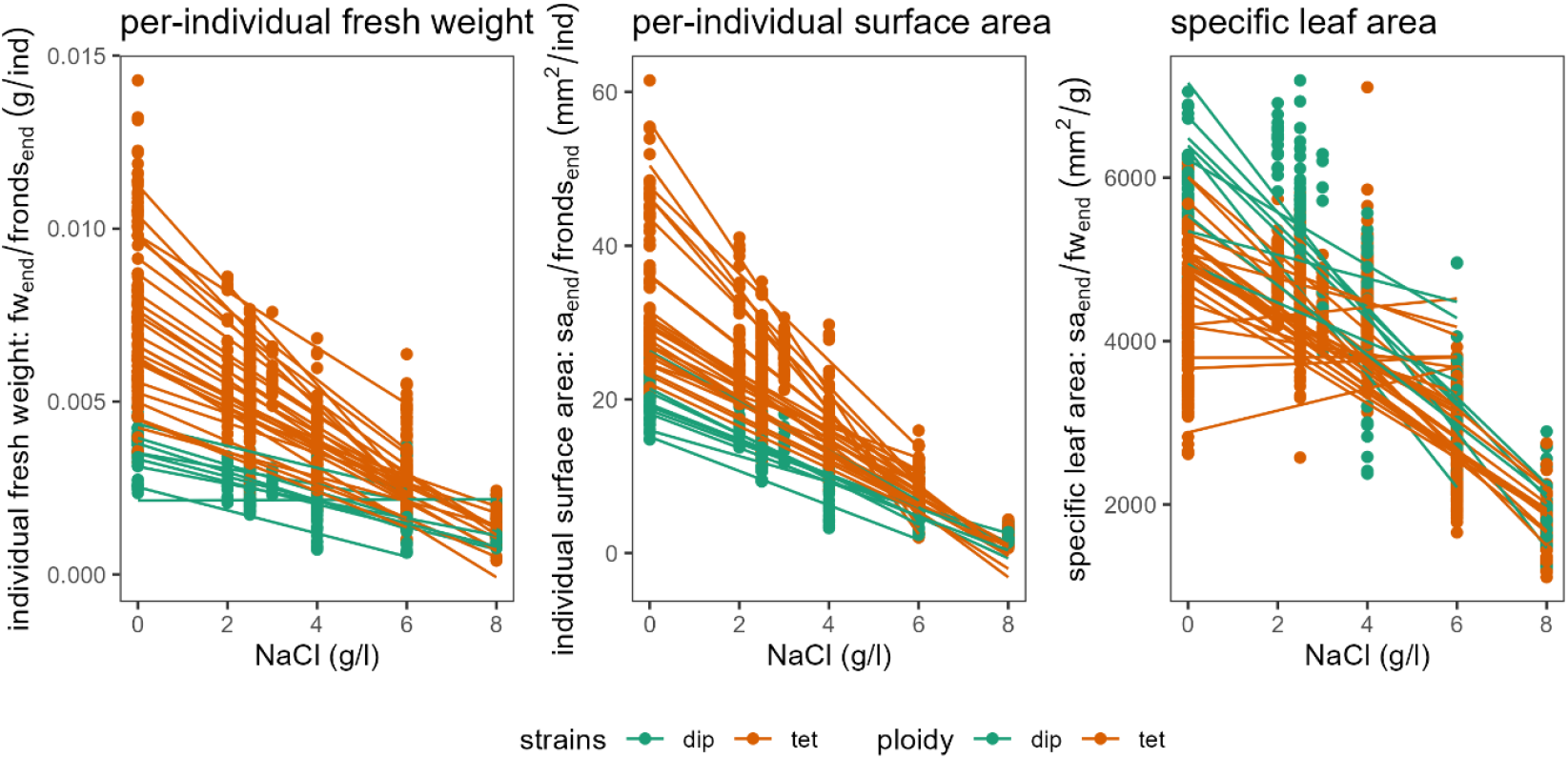
average morphological traits across the salt gradient: 1) average individual fresh weight, 2) average frond surface area and 3) average fresh weight specific leaf area across three independent colchitetraploid strains (red) compared to their progenitor diploid strain (green) for nine different genotypic backgrounds. Trend line of best fitting linear regression per clone to indicate direction of change across salt gradient.

## Discussion

Our experiment provides an explicit, replicated test of how much phenotypic variation WGD per se induces relative to genotypic sequence diversity and environment. Across nine genotypes, three independent autotetraploids per genotype, and a salt gradient, deterministic effects of ploidy explain more variation in growth and reaction norms than all genetic sequence variation combined, with limited-yet-unmistakable stochasticity as demonstrated by rare outliers.

In benign conditions, diploids tend to grow faster, whereas tetraploids exhibit larger, thicker fronds, confirming earlier work (Bafort *et al*., 2023; Anneberg *et al*., 2023). Under salt stress, the relative performance of tetraploids converges and often overtakes that of the diploid progenitor, indicating a predictable shift in phenotype after genome doubling: reduced growth at low stress, enhanced tolerance at high stress.

Among 26 tetraploid strains, only one strain, 9500c, displayed a similar relative growth rate compared to its progenitor diploid. Another tetraploid strain, 9502b, displayed an exceptionally low growth rate compared to the other tetraploids of genotype 9502, indicating the importance of rare stochastic effects during WGD. However, we cannot rule out mutations during the generations (ca. 50) between polyploid induction and growth testing. The diploid advantage in control conditions diminishes with increasing salt concentration to the point that most tetraploid strains recorded an equal and some even higher growth rate compared to their diploid progenitor strain. This confirms earlier-reported salt-tolerance in tetraploid *Spirodela polyrhiza* (Bafort *et al*., 2023), and is associated with bigger individuals in terms of biomass and surface area and with a lower specific leaf area.

WGD had a clear non-stochastic effect on growth rate. Indeed, growth rates among tetraploids of the same genotypic background (i.e., descending from the same diploid strain) overall differed very little compared to growth rates between genotypes and cytotypes. The aberrant growth rate of tetraploid 9500c, matching the growth rate of its diploid progenitor, shows that performance deviations are rare but possible. However, this unique diploid-like tetraploid strain 9500c did not exhibit the same sensitivity to salt as the diploid. Therefore, WGD showed an even more consistent effect on salt tolerance. Aside from this outlier, the consistent effect of polyploidy across our duckweed panel is striking compared to the high variability within genotypes reported by Aversano et al. (2013).

We further showed that the overall effect of the salt gradient, so the environmental component of phenotypic expression, was the largest source of variation in growth rate across our experiment. This is not surprising given that duckweeds are freshwater organisms and the fact that we choose the salt gradient to include extreme, though viable, levels. We found that both the deterministic effect of polyploidy and differences in genotypic background explained a substantial share of variation, exceeding the residual variation under control conditions. The deterministic ploidy effect outweighed the influence of genotypic backgrounds on growth, both in the control conditions and in terms of tolerance to salt, with the genotype-specific ploidy effects explaining only a smaller part of variation. Note that the nine genotypic backgrounds we used encompasses the global distribution and phylogenetic diversity of *Spirodela polyrhiza*, whose total genetic variation has been deemed low (Xu *et al*., 2019). This suggests that, in greater duckweed, polyploidy has a stronger impact on growth than the combined effects of genomic sequence diversity, which traditionally accounts for the genetic component of phenotypic heterogeneity.

In the control conditions, higher growth rate was associated with strains that have lighter and smaller fronds with a higher specific leaf area (i.e., thinner fronds). The fast-growing tetraploid 9500c likewise showed this “diploid” morphology. The strong association indicates that the larger size and slower growth rate have a common underlying cause, such as a larger cell size. Whatever the underlying cause, the predisposition to develop bigger but thicker fronds may entail an innate ability for polyploids to implement the common salt stress response to thicker (but smaller) leaves (Munns and Tester, 2008) as a side effect of WGD. We, indeed, observed a decreasing specific leaf area as the common response to high concentrations of salt in all strains, which reduces the area of contact of these surface floating organisms with the saline medium per unit of photosynthetic tissue and may hint at larger vacuoles to offload accumulating Na^+^ ions out of the cytoplasm. However, this does not explain how the diploid-like tetraploid 9500c still tolerated the salt stress conditions equally well compared to other tetraploids. While it is possible that the more robust polyploid fronds buffered the salt response better, the morphological consequence of polyploidy likely did not increase salt tolerance directly.

The rather small influence of stochastic processes that we found on the polyploid phenotype may stem from its asexual reproductions that avoids any mutational effects recombination. This means that the consequence of WGD may be inherently more stochastic in sexual systems compared to asexual systems. At the same time, it is hard to uncover such stochastic consequences in sexual polyploid systems. The reshuffling effect of meiosis in each generation results in independent genome duplication events occurring in differently shuffled genotypic backgrounds, even when closely related individuals produce polyploids independently. Generating independent polyploids from inbred lines could shed light on the severity of stochastic consequences in sexual polyploids.

There are good indications that WGD can trigger molecular mechanisms that lead to stochastic, or unpredictable, phenotypic outcomes. A well-studied phenomenon is “genomic shock”, a process primarily associated with allopolyploids that encompasses large structural mutations shortly after polyploid formation due to differentiation between subgenomes (Comai *et al*., 2003; Shimizu, 2022; de Tomás and Vicient, 2024). Whether genome doubling alone can trigger stochastic processes similar to those seen in polyploid hybrids remains uncertain. Autotetraploids rarely exhibit the extensive structural rearrangements tolerated in allopolyploids, though evidence is limited (Parisod, Holderegger and Brochmann, 2010). Stable polyploids often show a doubled but otherwise unchanged genome (Ozkan, Tuna and Galbraith, 2006; Aversano *et al*., 2013). Still, there is ample evidence for polyploidy-caused transposon activity (Bardil, Tayalé and Parisod, 2015; Vicient and Casacuberta, 2017; Baduel *et al*., 2019; He *et al*., 2022), epigenetic changes (Yu *et al*., 2010; Aversano *et al*., 2013), and changes in nuclear architecture and nuclear surface area to volume ratio that both affect nuclear topology. When autopolyploidy affects gene expression (but see Allario *et al*., 2011), their strength and direction are highly gene-specific (Guo, Davis and Birchler, 1996; Coate and Doyle, 2010; Visger *et al*., 2019; Song *et al*., 2020), adding to the inherent “noisiness” of gene expression (Kempe *et al*., 2015; Araújo *et al*., 2017). Affected genes potentially contain a small subset involved in the cell’s response to a duplicated genome (Yu *et al*., 2010; Fasano *et al*., 2016) or in a concerted gene dosage regulation.

Our experiment shows that independent genome duplication events can have stochastic consequences. Despite this possibility, nearly all phenotypic heterogeneity attributable to genetic factors was explained by the deterministic effect of polyploidy and genotypic background. Of these, polyploidy exerted a stronger influence than the genomic sequence diversity, both as a genetic component and in its interaction with the salt gradient. We therefore conclude that WGD generally produces consistent stress tolerant phenotypic effects on morphology and fitness. Genotype-specific consequences immediately following WGD likely reflect strong, repeatable responses rather than random variation. Nonetheless, the fast-growing outlier tetraploid strain 9500c and slow-growing outlier 9502b demonstrate that rare stochastic deviations can occur.

## Material and methods

### Plants

We use greater duckweed (*Spirodela polyrhiza*) as a model system (Laird and Barks, 2018) to test the consequences of polyploidy. Duckweeds (Lemnoideae) form a subfamily of free-floating fresh water plants, being the smallest and fastest reproducing Angiosperms. They mainly reproduce clonally. We started from nine different diploid genotypes (0013, 0225, 9242, 9316, 9346, 9500, 9502, 9503, 9512, *supplementary materials 1* for more information) and developed three colchitetraploid strains from each diploid genotype (cfr. Wu *et al*., 2023). The three tetraploid strain for each genotype, therefore, signify repeated polyploidy events and will be referred to as replicated tetraploids (or tetraploid strains). Figure 1 show the structure in tested strains and therefore the sources of variability that affect measurements of this collection of strains.

In short, we applied a colchicine treatment (Hoagland’s E medium with 0.7% colchicine and 0.5% DMSO for 24h; details in Bafort et al., 2023) to single fronds (used here to refer to single individuals, consisting of a single thallus with leaf and rhyzomes). We confirmed ploidy on surviving fronds using flow-cytometry and selected three stable tetraploid strains per progenitor diploid genotype. We did not obtain diploid strains that underwent the same colchicine treatment to control for other possible mutagenic effects of colchicine and DMSO. While expected to be low, we cannot completely rule out variation generated between independent colchitetraploid strains because of these compounds.

### Population growth tests

We tested all strains in Hoagland’s E medium (Okunowo and Ogunkanmi, 2010) supplemented with 0, 2.5, 4 and 6 g/l NaCl and tested 8g/l NaCl in all strains of genotype 0225, 9242, 9500 and 9503. Prior to testing, all strains were pretreated for two weeks under identical laboratory conditions, using the same type of containers and the same type of medium as in ensuing growth tests. For each test, we weighed 20 fronds across all life stages that were gently dry patted to determine fresh weight. These fronds were then added to 100ml of the correct medium in a transparent plastic box (1l) covered with square petri dish lid that allows sufficient air circulation. Each box was photographed (without lid) in a standardized setup, and the number of starting individuals was recorded to correct for potential counting mistakes in the previous step. After 7 days, we counted all fronds (again, across all life stages), repeated the fresh weight measurements and photographs and added the plant material to pre-weighed aluminum foil envelopes to dry in a ventilated oven at 70°C for 5 days to measure dry weight. Pilot tests confirmed that this temperature and duration was sufficient to have all plant moisture evaporate. None of the tested replicates produced sinking reduced fronds that function as resting propagules, called turions. We observed reduced fronds at higher salt concentrations that were visually different from turions and were also not expected to be since salt inhibits turion formation (Kuehdorf and Appenroth, 2012). Photographs were taken with a gray scale video camera and processed in ImageJ to calculate total frond surface area using the “Analyze particles” implementation to detect groups of fronds. We adjusted the greyscale threshold values per batch to account for minor differences in light conditions, ensuring accurate detection of all green frond tissue while excluding background.

Growth experiments were conducted in multiple batches from November 2023 to December 2024. Each batch included all derived strains (diploid and tetraploid) from at least two progenitor genotypes tested under a certain salt treatment. In cases where populations failed to grow sufficiently during pretreatment, they were excluded from the corresponding batch.

### Ploidy verification with flow cytometry

At the end of the experimental period, we verified ploidy for all 36 strains used in this study using the protocol found in (Wu *et al*., 2023). We found that all strains exhibited the expected ploidy except for 9512b that was found to be mixoploid, containing a fixed proportion of diploid and tetraploid cells. We show its growth rate but omit this strain from further analyses in this study. More details on protocol and results can be found in supplementary materials (Appendix 4)

### Statistics and morphological traits

We analyzed growth in the control medium by modelling the natural logarithm (i.e., ln) of either count, fresh weight (*fw*) or frond surface area (*sa*) at day 7 as a response with normal error distribution of the log count, *fw, sa* on day 0 added with a relative growth rate (*rgr*). *Rgr* is affected by a linear combination of effects from genotype (*geno*), ploidy, their interaction and a fixed effect for each independent tetraploid strain within each genotypic background to model deviations in *RGR* between independent tetraploid strains of the same progenitor genotype (*geno:strainl*).

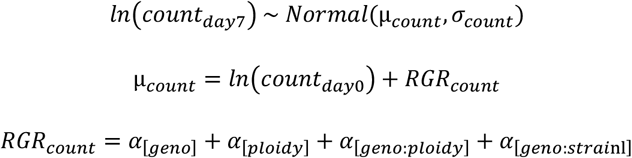

This assumes an exponential growth in numbers, biomass and surface area. This is applicable for our duckweed populations because we put them in low-competition conditions of 20 fronds in a relatively large space growing over a relatively short period. The response variables (*count*_*day*7_, *fw*_*day*7_, *sa*_*day*7_) were modelled as correlated.

We extended the statistical model to infer effects of genotype (*geno*), ploidy and their interactions across the salt gradient (B). This reaction norm to salt represents the environmental component of the measured heterogeneity in growth and infers how genetic factors affect this reaction norm (i.e., GxE interactions). We also model a replicate tetraploid strain variable effect (*geno:strainl*) on the salt slope. We again model the response variables (*count*_*day*7_, *fw*_*day*7_, *sa*_*day*7_) as correlated. For *RGR* of *fresh weight*, we formulate the model extension from the previous as follows:

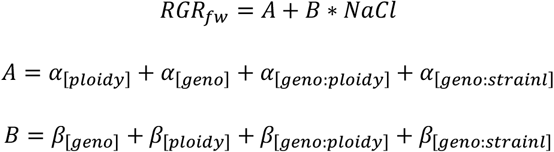

We elected to model the effect independent tetraploid strain within each genotype (*α*_[*geno:strainl*]_, containing the stochastic effect of polyploidy) in both models as a fixed rather than as sampled from a distribution of possible independent polyploid strains (i.e., in a mixed-effects model). In a mixed-effects model, we would estimate an inaccurate group-level standard deviation from only three independent tetraploid strains per genotypic background. Categorical variables for the genotypic background (*geno*) and independent tetraploid strain (*geno:strainl*) are centered to estimate the other effects on the overall mean across genotypes and tetraploid strains within genotype rather than conditional on one arbitrary reference genotype or tetraploid strains, respectively (Schielzeth, 2010). The independent tetraploid strain variable is centered to the mean within each genotypic background. We did not center the *ploidy* variable as we deem the diploid as an appropriate reference level to estimate differences between genotypes, independent tetraploid strains and applied salt concentrations.

We partitioned the different sources of variability in these models by calculating the (finite-sample) standard deviation of estimated coefficients within each effect. We estimate a genotype-specific standard deviation among independent tetraploid strains, to consider between genotype differences in stochastic WGD effects. In the salt gradient model, we scale the standard deviation of the slope across the salt gradient, and all its interactions, to the response by multiplying said standard deviation with the standard deviation in recorded salt concentrations (Gelman and Hill, 2006).

From counts and measurements, we calculated population-averaged traits such as individual fresh weight 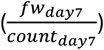, individual surface area 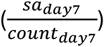, fresh weight-based leaf specific area 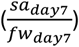 and dry matter proportion 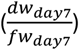. All metrics except dry weight proportion can also be estimated using the estimated correlations from the statistical models.

All analyses were performed in R (R core team, 2021) using the brms library (Bürkner, 2018) for Bayesian statistics, based on Stan (Stan Development Team, 2019), and the tidyverse ecosystem (Wickham *et al*., 2019). Bayesian inference was performed using Hamiltonian Monte Carlo (HMC) models that implemented four chains with each 4,000 iterations from which 2,000 were considered burn-in. We evaluated the performance of every fitted model based on standardized procedures by checking mixing and stationarity in the trace plots and by checking the effective sample size and 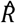 statistic for each parameter (McElreath, 2020). Models, formulae, and detailed description of the procedure and priors can be found in the electronic supplementary material.

### Dry weight

Dry weight is tightly correlated with fresh weight within each genotype. Therefore, we assume that relationships between fresh weight growth and other responses are transposable to dry weight growth. More information in supplementary materials (Appendix 13, Figure S13).

## Acknowledgements

We thank Annelore Natran and Jesse Terryn for stock culture maintenance and help during the creation of colchitetraploid clones, Silvija Molisavljevic for providing the global map with locations of origins of the diploid clones, and dr. Maxime Dahirel for advice on the statistical modelling. F.M. acknowledges funding from Fonds Wetenschappelijk Onderzoek (FWO, 1238124N). Y.V.d.P. acknowledges funding from the European Research Council (ERC) under the European Union’s Horizon research and innovation program (No. 833522) and from Ghent University (Methusalem funding, BOF.MET.2021.0005.01).

## Conflict of interests

No conflict of interests

### Data availability statement

Data and R code to analyse the data and generate figures are available at: https://github.com/fremorti/polyploid-repeatability/ ; and will be published to zenodo (together with flow cytometry files) at moment of article publication.

