## Appendix for "Highly repeatable phenotypic consequences of whole-genome duplication in *Spirodela polyrhiza*"

#### Appendix 1: Spirodela polyrhiza diploid genotypes

We started our panel of 36 strains to test from nine diploid genotypes: 0013, 0225, 9242, 9316, 9346, 9500, 9502, 9503, 9512. We received these lines from the kind folks at the Landolt collection in Zürich, which has since been moved. These genotypes represent the global distribution of origins of greater duckweed (fig. S1) and harbors members of all four population genetic clusters of *S. Polyrhiza*, which has been measured as quite low (Xu et al., 2019).

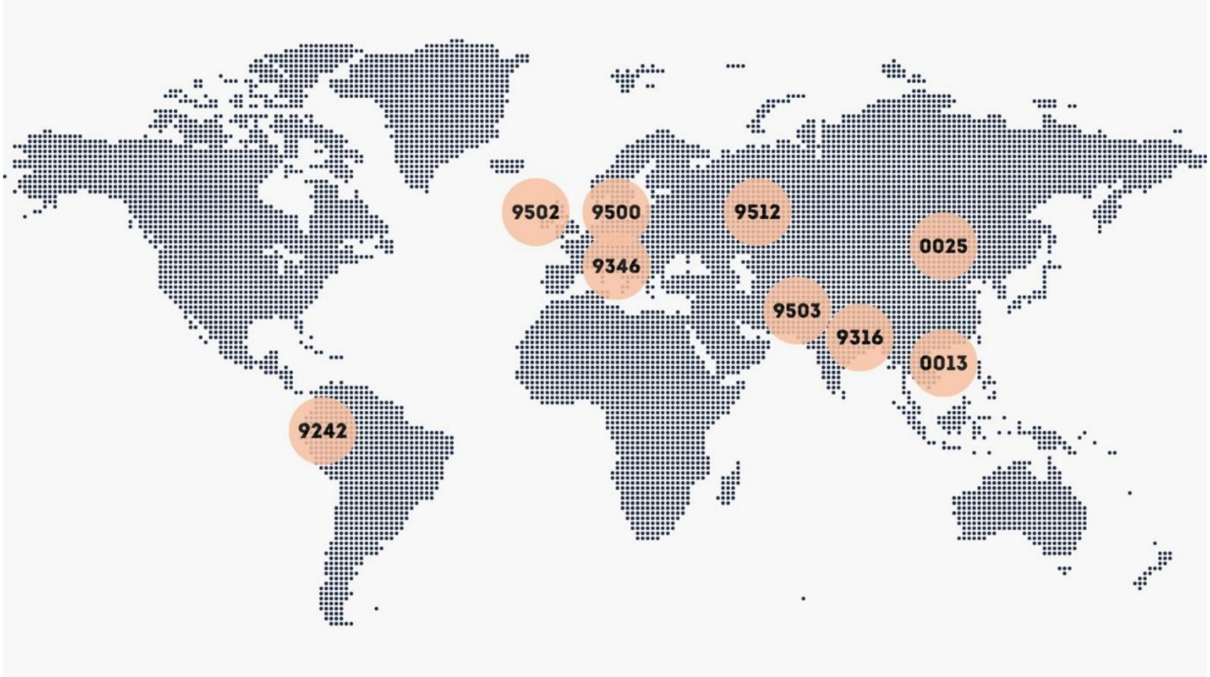


*Figure S1: Global map of locations from which the genotypes were originally sampled. Map created by Silvija Milosavljevic*

#### Appendix 2: complete model formulation of growth in control medium

$$\begin{bmatrix} \ln(count_{day7}) \\ \ln(fw_{day7}) \\ \ln(sa_{day7}) \end{bmatrix} \sim MVNormal(\begin{bmatrix} \mu_{count} \\ \mu_{fw} \\ \mu_{sa} \end{bmatrix}, \begin{bmatrix} \sigma_{count}^2 & \rho_{count,fw} * \sigma_{count} * \sigma_{fw} & \rho_{count,sa} * \sigma_{count} * \sigma_{sa} \\ \rho_{count,fw} * \sigma_{count} * \sigma_{fw} & \sigma_{fw}^2 & \rho_{fw,sa} * \sigma_{fw} * \sigma_{sa} \\ \rho_{count,sa} * \sigma_{count} * \sigma_{sa} & \rho_{fw,sa} * \sigma_{fw} * \sigma_{sa} & \sigma_{sa}^2 \end{bmatrix})$$

$$\mu_{count} = \ln(count_{day0}) + RGR_{count}$$

$$\mu_{fw} = \ln(fw_{day0}) + RGR_{fw}$$

$$\mu_{sa} = \ln(sa_{day0}) + RGR_{sa}$$

$$RGR_{count} = \alpha_{count,[geno]} + \alpha_{count,[ploidy]} + \alpha_{count,[geno:ploidy]} + \alpha_{count,[strainl]}$$

$$RGR_{fw} = \alpha_{fw,[geno]} + \alpha_{fw,[ploidy]} + \alpha_{fw,[geno:ploidy]} + \alpha_{fw,[strainl]}$$

$$RGR_{sa} = \alpha_{sa,[geno]} + \alpha_{sa,[ploidy]} + \alpha_{sa,[geno:ploidy]} + \alpha_{sa,[strainl]}$$

Coefficients $\alpha_{metric,[geno]}$, $\alpha_{metric,[geno:ploidy]}$ and $\alpha_{metric,[strainl]}$ are each a linear combination of effects of centered dummy variables and their estimated coefficients. There is a centered dummy variable for each level minus one of genotype (8 dummy variables, *$is0225_c$, $is9242_c$, $is9316_c$, $is9346_c$, $is9500_c$, $is9502_c$, $is9503_c$*, and *$is9512_c$*) and of independent tetraploid strain (2 dummy variables, $isa_c$ and $isb_c$) according to (Schielzeth, 2010). To center each dummy variable, the dummy value (so, 0 or 1) is subtracted by the mean of that dummy variable, equalling the frequency of the level the dummy variable represents. In the case of the *strainl* dummy variables, the value for each diploid is put at 0 and the centering for each tetraploid strain is done within each genotype, not considering the diploid of that genotype, so that the independent polyploid strain effect is modelled as a deviation from that genotypes mean (deterministic) polyploid effect. As an example, $\alpha_{count,[strainl]} = \alpha_{count,isa} * isa_c + \alpha_{count,isb} * isb_c$, with $isa_c$ and $isb_c$ the centered dummy variables for independent tetraploid strain a and within each genotype, respectively.

#### Appendix 3: model formulation of greowth in salt gradient extension

We extend the model of growth in control medium (Appendix 2) with for each metric (i, either count, fw or sa) an intercept and slope across the salt (NaCl) gradient that is modelled with genotype (geno), ploidy, their interaction and a variable effect for each replicated polyploid strain (strainl).

$$RGR_i = A_i + B_i * NaCl$$

$$A_i = \alpha_{i,[geno]} + \alpha_{i,[ploidy]} + \alpha_{i,[geno:ploidy]} + \alpha_{i,[strain_l]}$$

$$B_i = \beta_{i,[geno]} + \beta_{i,[ploidy]} + \beta_{i,[geno:ploidy]} + \beta_{i,[strainl]}$$

The effects involving *geno* or *strainl* are, again, a linear combination of effects using centered dummy variables as explained in Appendix 2.

#### Appendix 4: flow-cytometry protocol and results

We used a nuclei extraction buffer containing 45mM MgCl2 + 6H2O, 30mM Sodium citrate + 2H2O, 20mM 4-morpholinepropane sulfonate (MOPS), 0.1% (vol/vol) Triton X-100 and 1% (mass/vol) Polyvinylpyrrolidone (PVP, Wu et al., 2023). This constitutes the buffer proposed by Galbraith and colleagues (1983), supplemented with PVP for binding cytosol phenolic compounds. In short, we sampled a single frond of each strain, chopped it with a razor in 0.5 ml nuclei extraction buffer to a homogenous suspension, filtered the resulting buffer with nuclei through a 40 µm mesh strainer and stained it with DAPI at a final concentration of 4 µg/ml. We processed each sample using an Attune NxT Acoustic Focusing Cytometer.

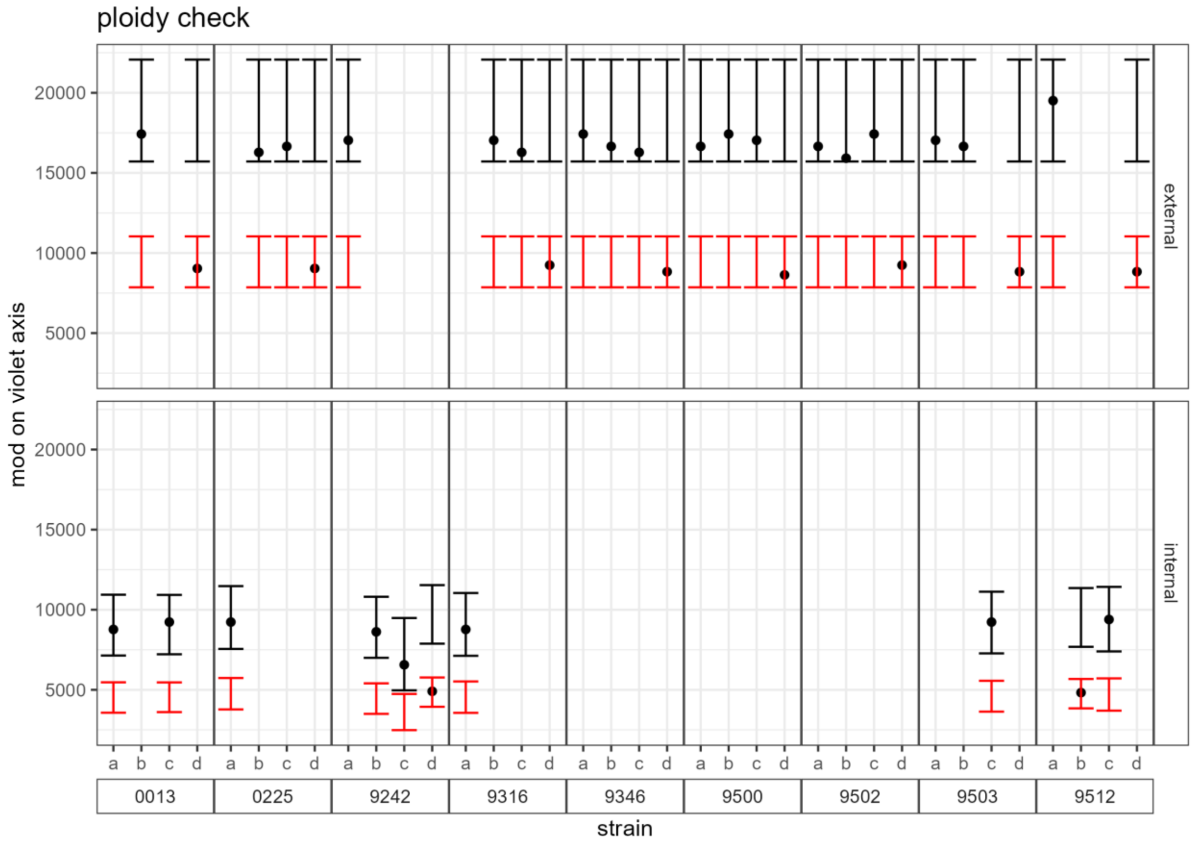


*Figure S2: ploidy check of all strains by comparing the measured modus on the violet (DAPI measuring) axis of the strain's nuclei in flow cytometric samples compared to an expected diploid range (red error bars) determined from a known diploid strain and an expected tetraploid range (black error bars) determined as the doubled lower and upper range limit of the expected diploid range. The samples with low quality results, bimodal distribution of nuclei or a mode outside the expected ranges in the first run with external control (upper panel) were repeated in a second ru with an internal control. We only show the second run results for those.*

Nuclear DNA content distributions were compared to that of an external sample of a known diploid *Spirodela polyrhiza* strain, measured in the same run as the other samples. The expected range of values on the violet axis (the sensor in the cytometer that measures the DAPI) for diploids was determined between the 1st and 99th percentile of the manually selected population of nuclei in the known diploid sample. The range for tetraploids was determined as double the value of the upper and lower range limit of the expected diploid nuclei range. Samples that showed low quality results (unclear nuclei population), lacked a clearly unimodal distribution of nuclei or showed a mode of their unimodal peak that fell outside the expected diploid or tetraploid range, were tested again by chopping in a diploid frond as an internal control. The mode of the biggest peak for the high-quality measurement falls within the expected range in all strains (Figure S2). For samples with internal control, which always had diploid peak of the control, we plotted the mode of nuclei distribution without the diploid nuclei population if the proportion of nuclei in the expected tetraploid range was higher than 0.25, so assuming more or less half the nuclei are from the diploid control. All strains had a unimodal nuclear DNA content distribution with their peak in the expected range, except for 9512b (Figure S3). This one strain had a bimodal distribution with a proportion of diploid nuclei and tetraploid nuclei. From further inspection, we determined that this proportion was relatively fixed across the population and across generations. Because of it mixoploid nature, 9512c was excluded from the analyses.

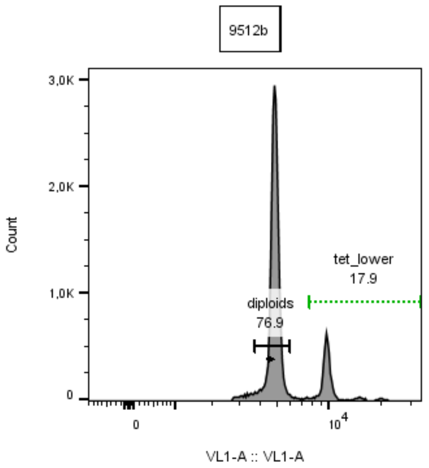


*Figure S3: DNA content distribution of potential nuclei from flow cytometry in strain 9512b. This strain had a bimodal distribution of diploid and tetraploid nuclei, indicating a mixoploid nature.*

### Appendix 5: relative growth rate in all metrics in control conditions

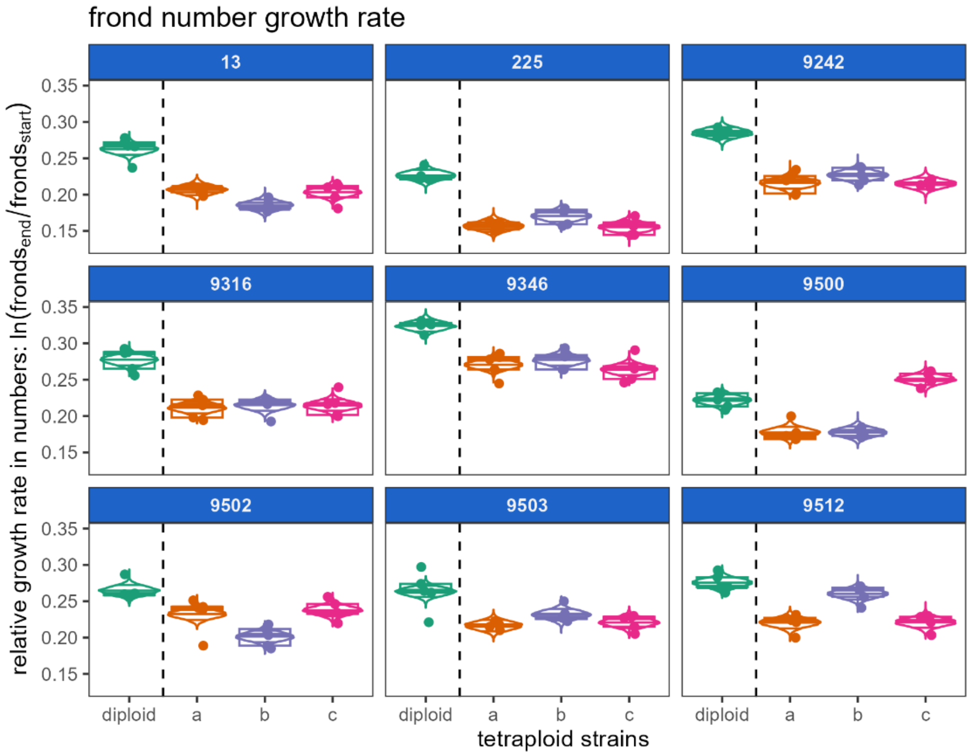


(1)

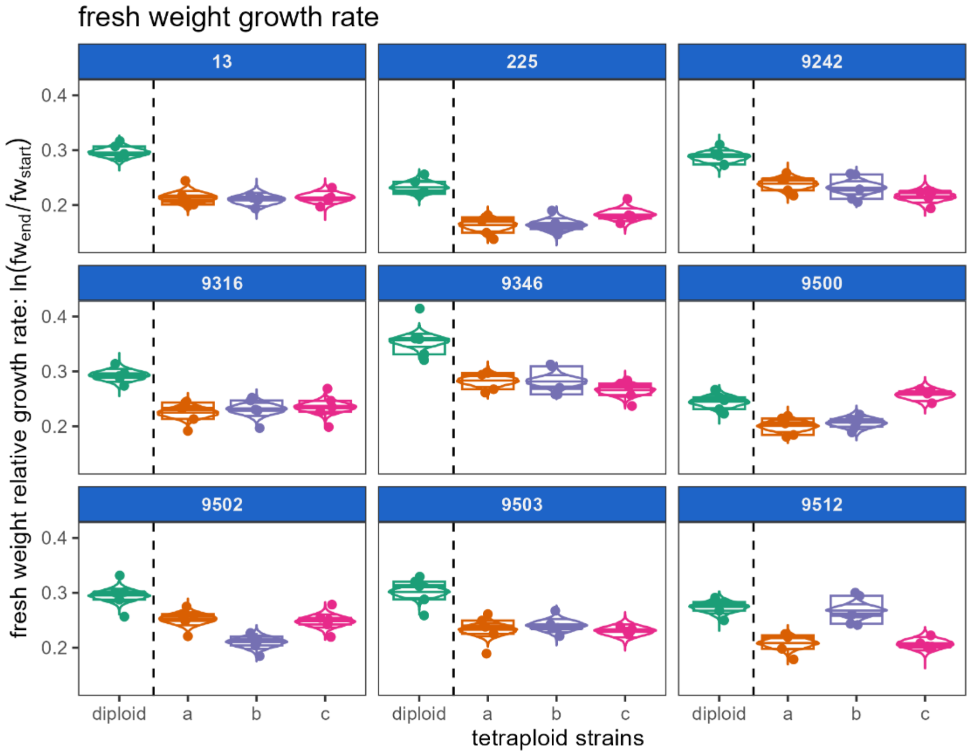


(2)

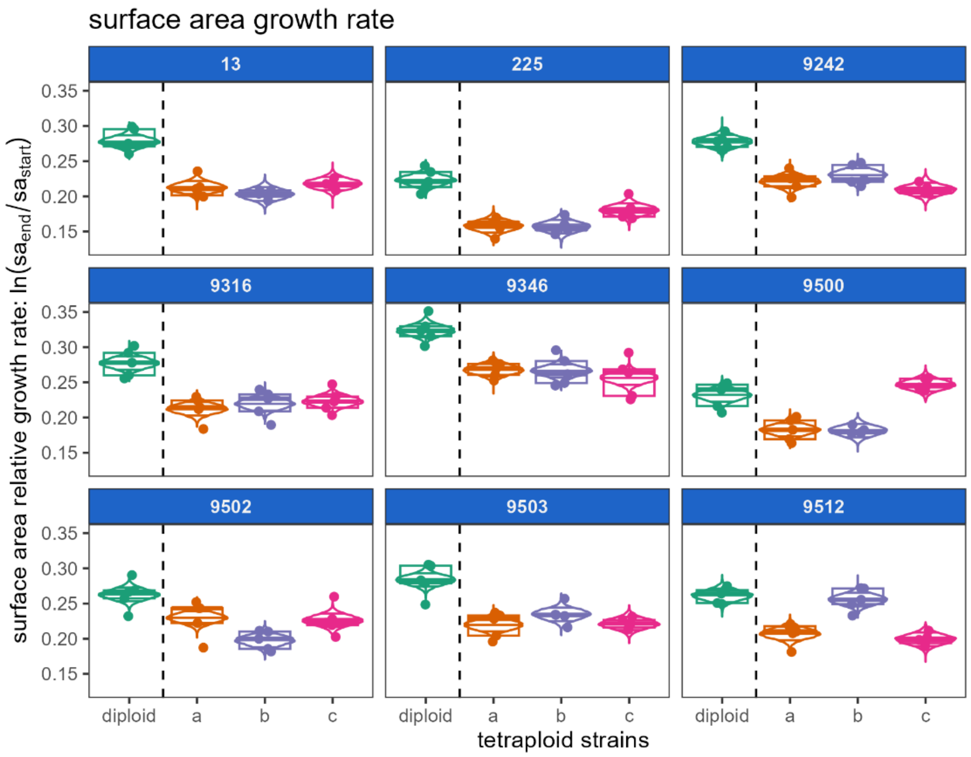


(3)

*Figure S4: relative growth rate (RGR) in control conditions in terms of number of fronds (1), fresh weight (fw, 2) and frond surface area (sa, 3) of three independent colchitetraploid strains (a, b, c) compared to their progenitor diploid strain for nine different genotypic backgrounds. Dots and boxplots show the observations and violins indicate the posterior expected RGR for that strain with the 9th, 50th and 91st percentile indicated. We show the results for the mixoploid strain 9512b, but exclude it for subsequent analyses.*

#### Appendix 6: variability by different factors for each growth metric in control conditions

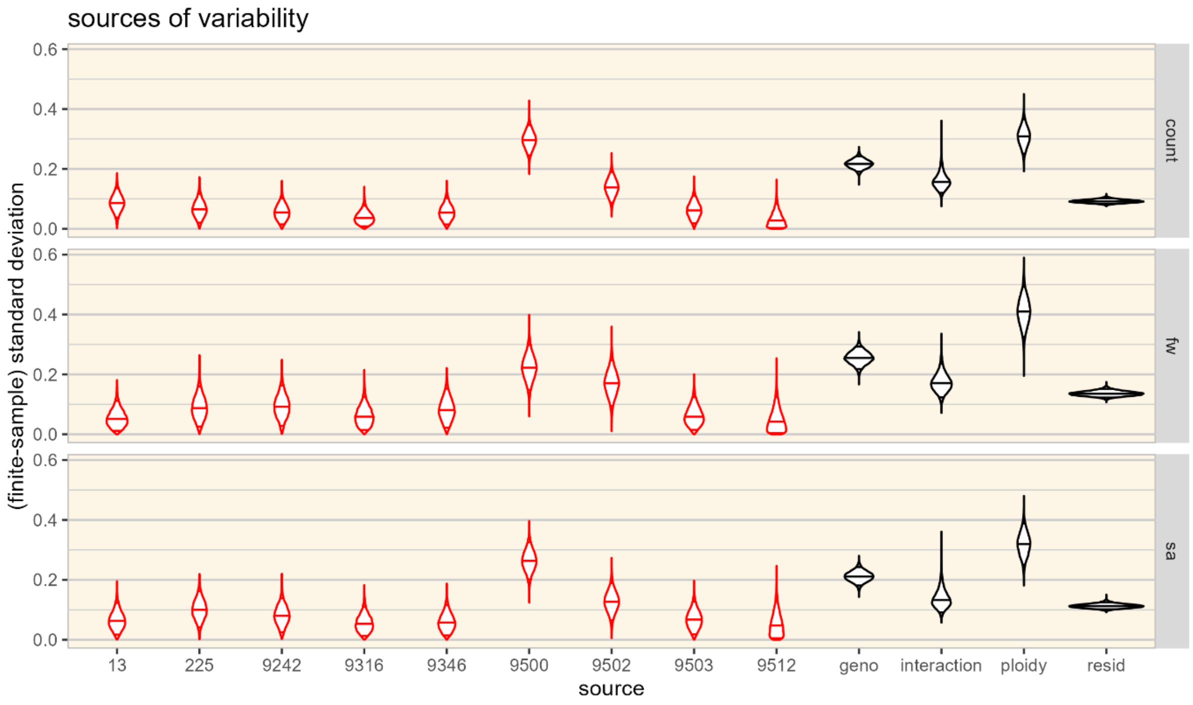


*Figure S5: posterior distribution of finite-sample standard deviation of all coefficients within each effect for the submodel of growth in count, fresh weight (fw) and frond surface area (sa) in the control condition. This shows the variation explained by independent polyploidy events (strain-specific, in red) compared to ploidy, genotype (geno), their interaction and the residual variation (black). We indicated the 4th, 50th and 96th percentile in each violin.*

### Appendix 7: correlation between growth in count, fresh weight and frond surface area

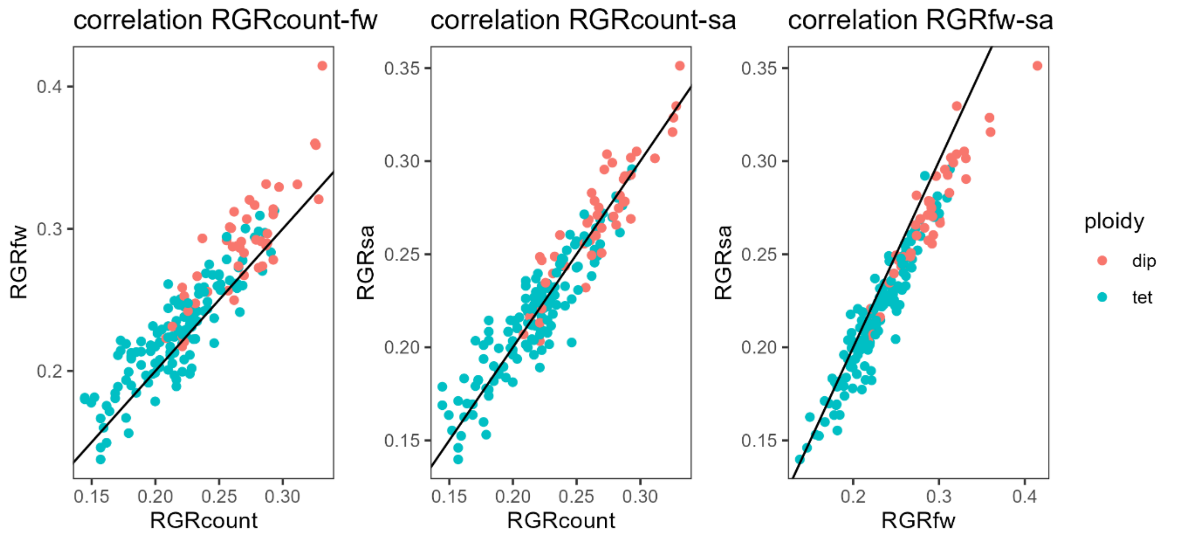


*Figure S6: correlation between growth in terms of count (RGRcount), fresh weight (RGRfw) and frond surface area (RGRsa) from data, where relative growth rate (RGRcount) was calculated as $ln\ (count_{day7}/count_{day0})$ and the others accordingly.*

### Appendix 8: correlations between morphology and growth rate in control conditions

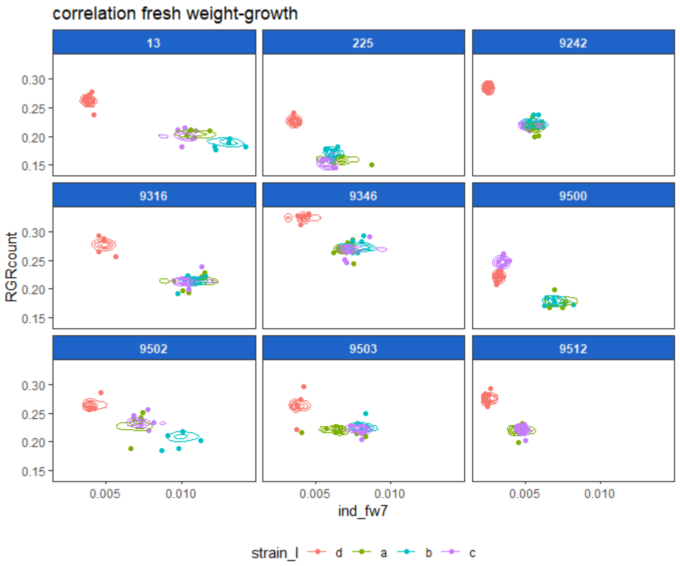


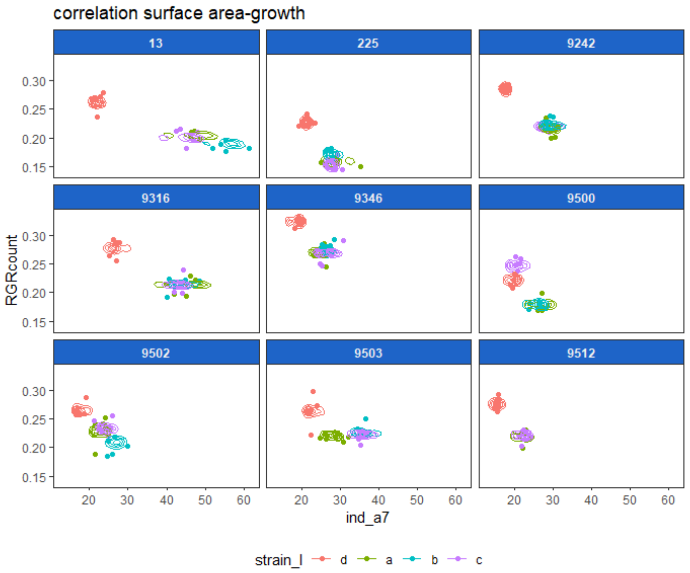

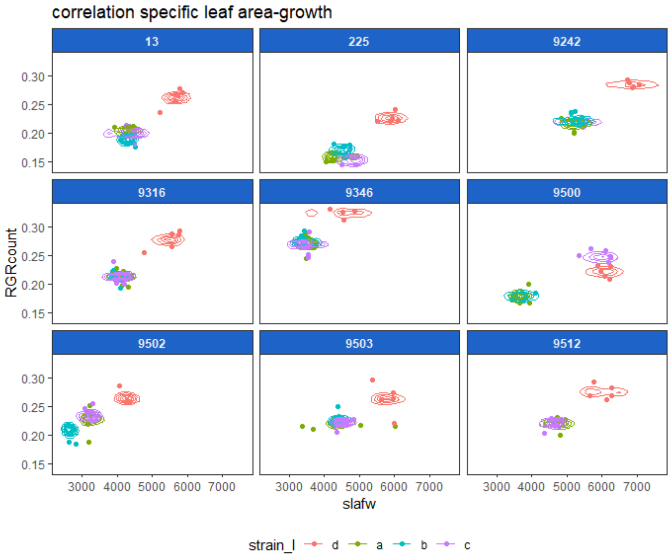


*Figure S7: correlations between growth rate in control conditions in terms of frond number count (1), fresh weight (2) and frond surface area (3) ) of three independent colchitetraploid strains (a, b, c) compared to their progenitor diploid strain (d, in red) for nine different genotypic backgrounds. Dots show correlations as calculated from data and concentric circles indicate areas of equal probability in the posterior distribution. We find strong correlations between strains of the same genotype bu no clear correlation within strains.*

### Appendix 9: relative growth rate at highest salt concentrations

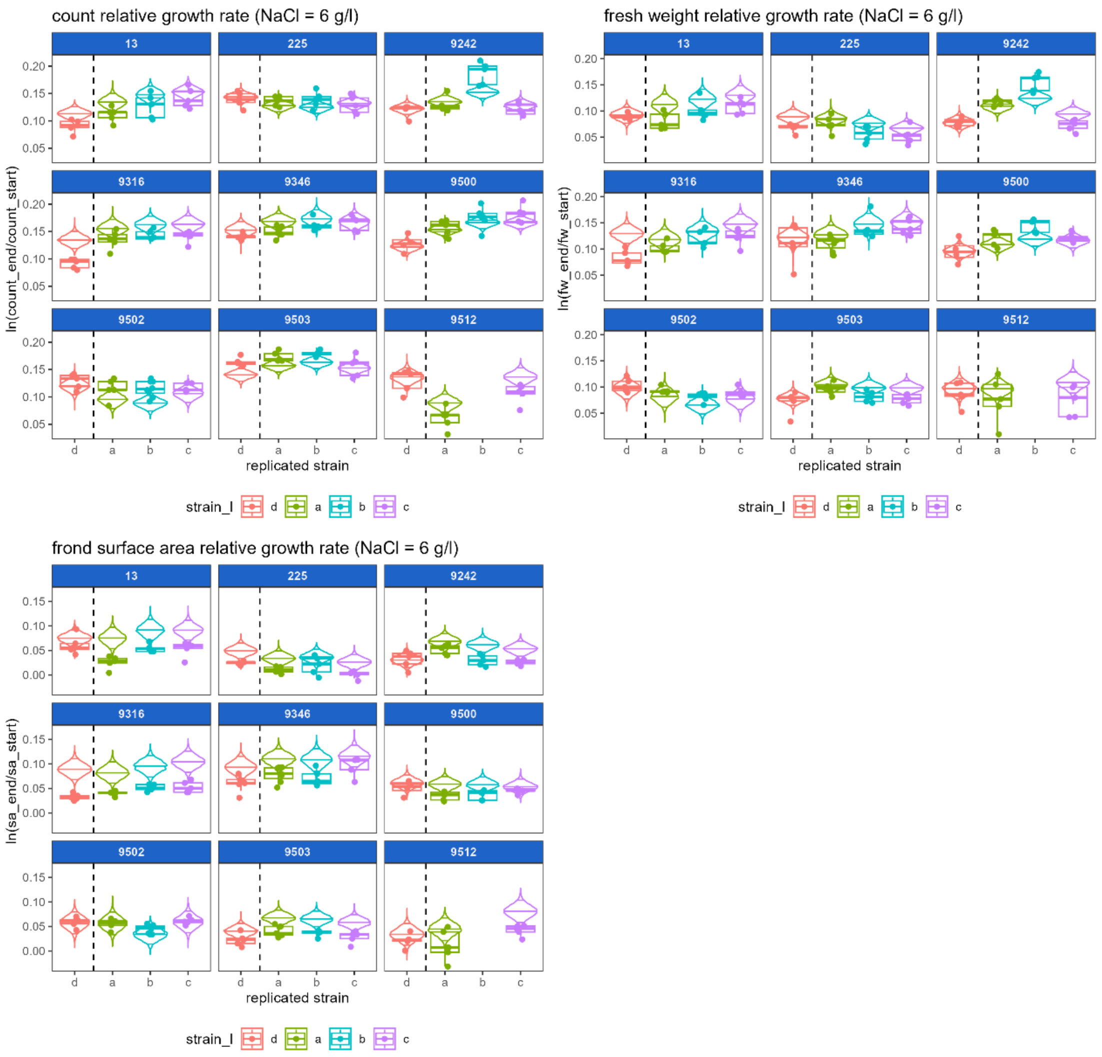


*Figure S8: relative growth rate (RGR) in 6g/l NaCl: in terms of count (1), fresh weight (fw, 2) and frond surface area (sa, 3) of three independent colchitetraploid strains (a, b, c) compared to their progenitor diploid strain (d, in red) for nine different genotypic backgrounds. Dots and boxplots show the observations and violins indicate the posterior expected RGR for that strain with the 4th, 50th and 96st percentile indicated. The expected RGR does not always fit well to the data because the fit is determined by the linear regression across the complete salt gradient where each concentration can deviate from this linearized decrease.*

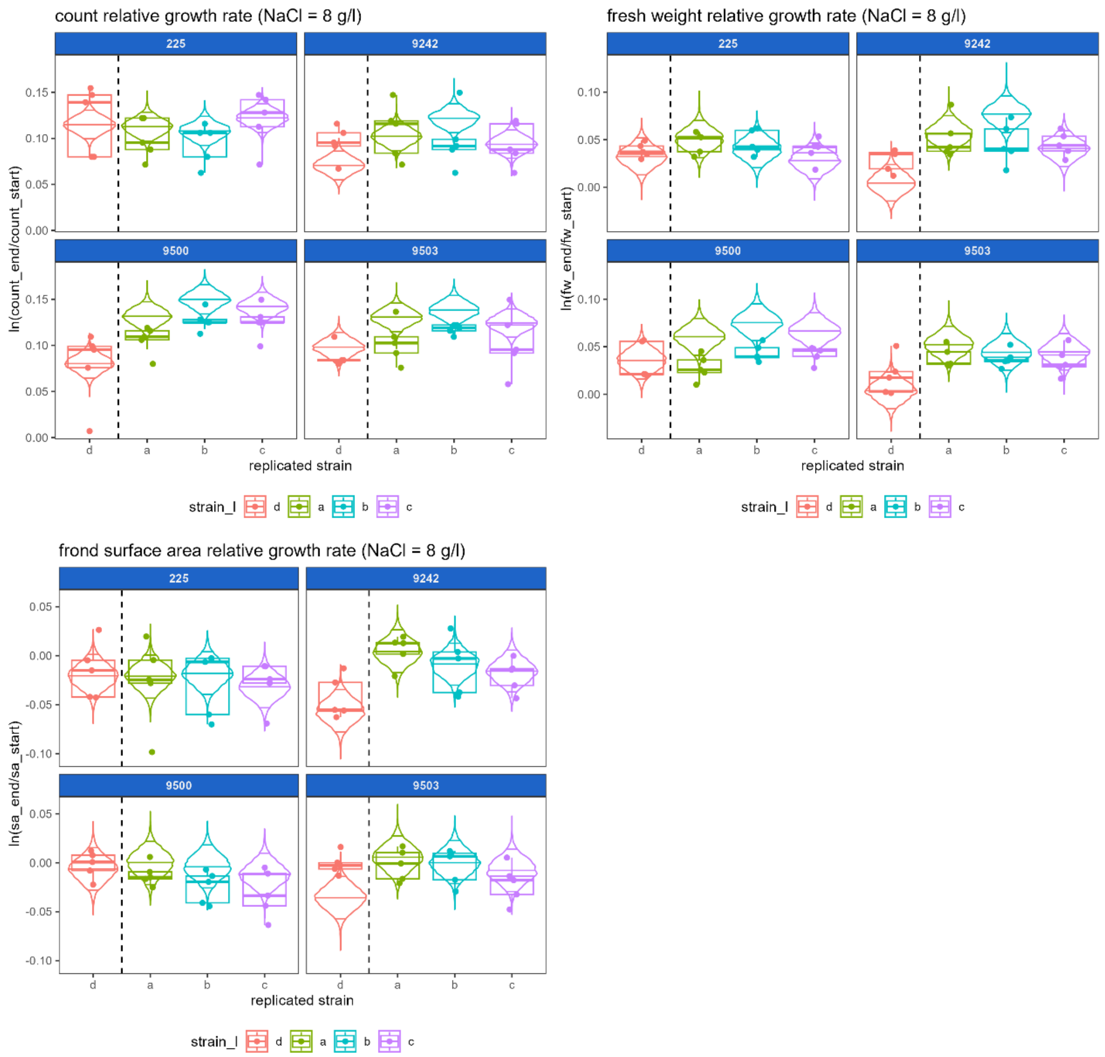


*Figure S9: relative growth rate (RGR) in 8g/l NaCl: in terms of count (1), fresh weight (fw, 2) and frond surface area (sa, 3) of three independent colchitetraploid strains (a, b, c) compared to their progenitor diploid strain (d, in red) for four different genotypic backgrounds. Dots and boxplots show the observations and violins indicate the posterior expected RGR for that strain with the 4th, 50th and 96st percentile indicated. The expected RGR does not always fit well to the data because the fit is determined by the linear regression across the complete salt gradient where each concentration can deviate from this linearized decrease.*

### Appendix 10: relative growth rate in all metrics across salt gradient

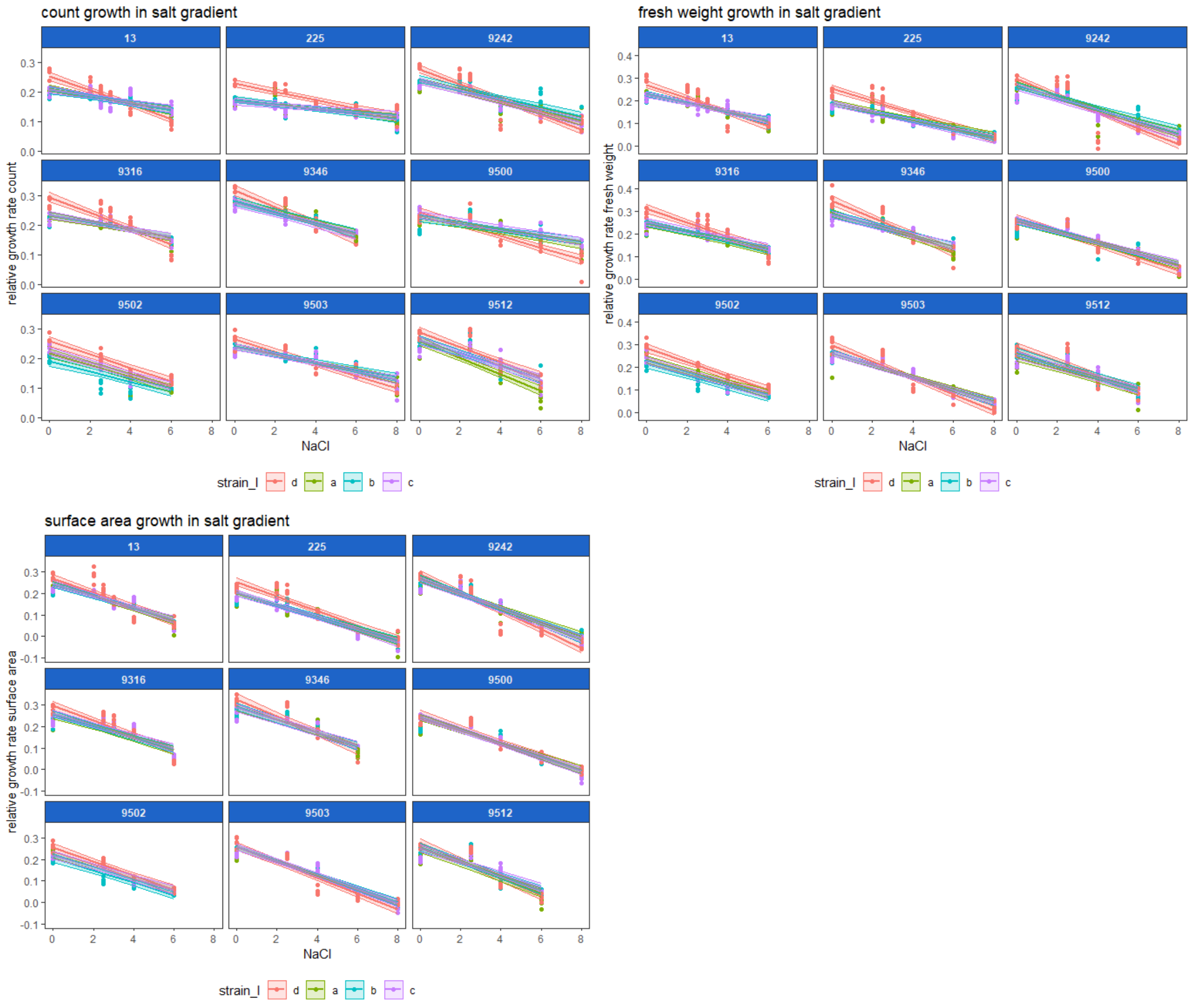


*Figure S10: Relative growth rate in count (1), fresh weight (fw) and frond surface area of three independent colchitetraploid strains (a, b, c) compared to their progenitor diploid strain (d, in red) for nine different genotypic backgrounds. Dots show the observations and lines indicate the median of the posterior expected RGR for that strain with the 4th and 96th percentile indicated by the ribbon.*

### Appendix 11: variability of intercept and slope across salt gradient of different factors for each growth metric

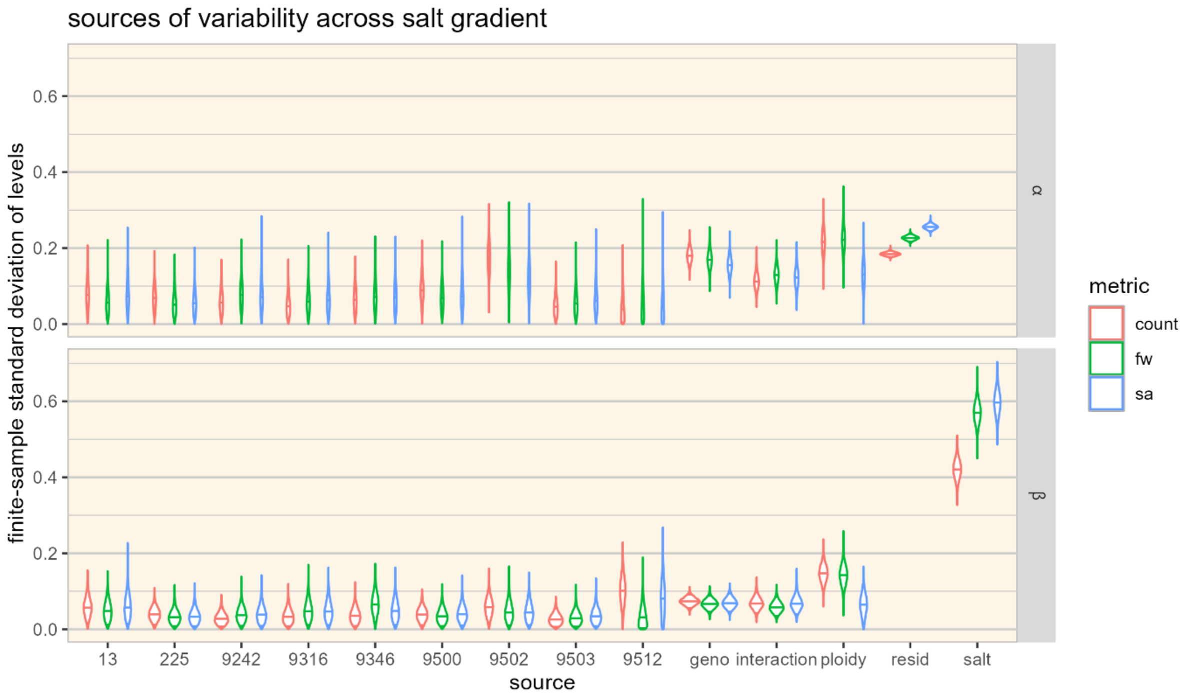


*Figure S11: posterior distribution of finite-sample standard deviation of all coefficients within each effect for the submodel of growth in count, fresh weight (fw) and frond surface area (sa) across the salt gradient. This shows the variation in intercept (α) and slope (β) explained by independent polyploidy events (strain-specific, in red) compared to ploidy, genotype (geno), their interaction and the residual variation (resid, black). The residual variation is on the same scale as the outcome and, therefore, only comparable with other sources of variation on the intercept. We indicated the 4th, 50th and 96st percentile in each violin.*

#### Appendix 12: correlations between morphology and growth rate across salt gradient

We found a negative correlation between individual weight and growth across strains in control conditions, but found a positive correlation across salt gradient. This means that our duckweed strains grow smaller individually and slower collectively with increasing salt concentrations.

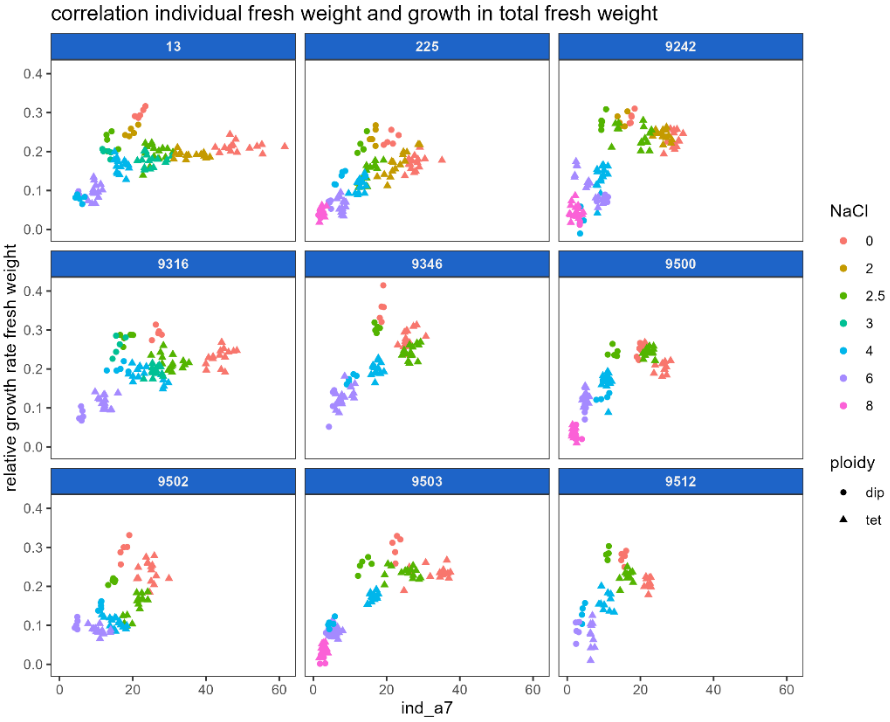
correlation individual fresh weight and growth in total fresh weight
13
225
9242
9316
9346
9500
9502
9503
9512
relative growth rate fresh weight
0.0
0.1
0.2
0.3
0.4
0
20
40
60
ind_a7
NaCl
0
2
2.5
3
4
6
8
ploidy
dip
tet

(a)

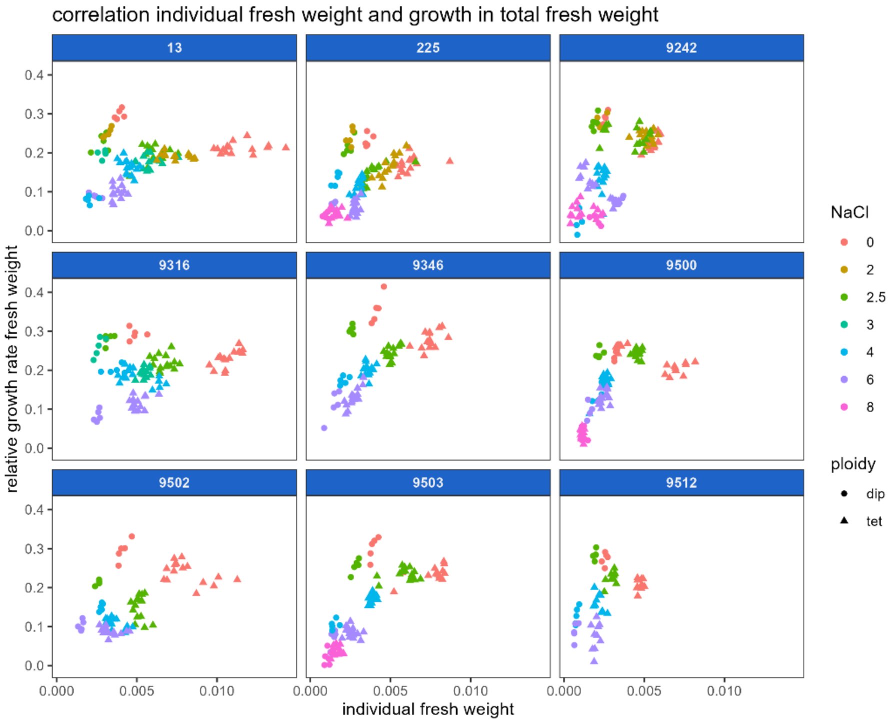
correlation individual fresh weight and growth in total fresh weight
13
225
9242
9316
9346
9500
9502
9503
9512
relative growth rate fresh weight
0.0
0.1
0.2
0.3
0.4
0.000
0.005
0.010
individual fresh weight
NaCl
0
2
2.5
3
4
6
8
ploidy
dip
tet

(b)

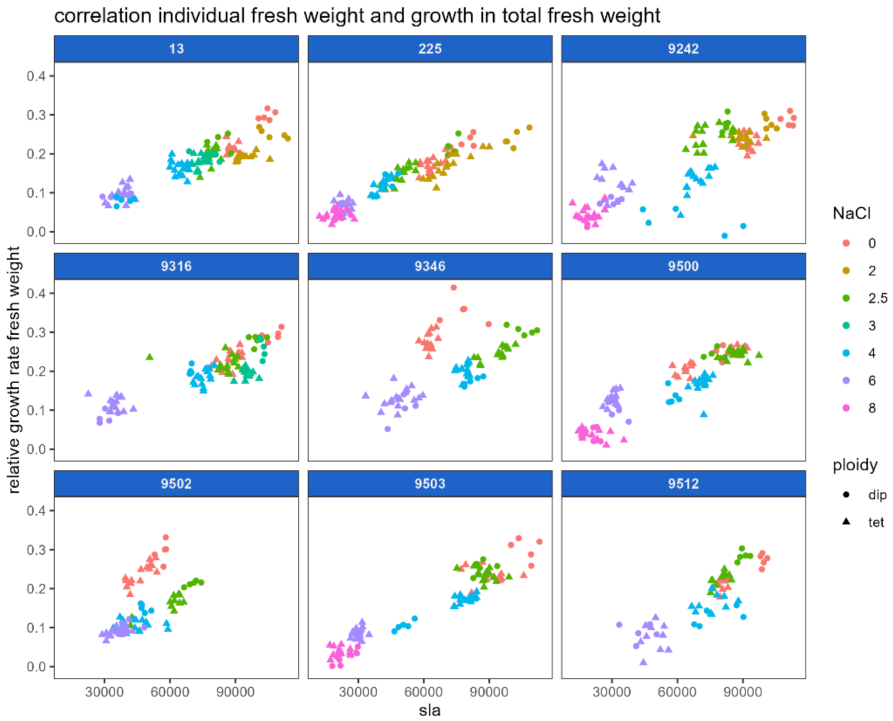


(c)

*Figure S12: relation between relative growth rate in fresh weight and individual fresh weight (a), individual surface area (b) and specific leaf area (c) across the salt gradient of three independent colchitetraploid strains (a, b, c, all in triangles) compared to their progenitor diploid strain (d, in dots) for nine different genotypic backgrounds.*

### Appendix 13: Dry weight

We present results from dry weight measurements separately since we were not able to model growth in dry weight (only a dry weight measurement at end of 7 day). Therefore, we cannot extract dry weight estimates from a posterior and can only show the recorded data.

Dry weight proportion is in some cases slightly higher in the diploid compared to the descendant tetraploids (in control, 9500, 9242, 9502) and increases slightly with increasing salt but is overall rather stable.

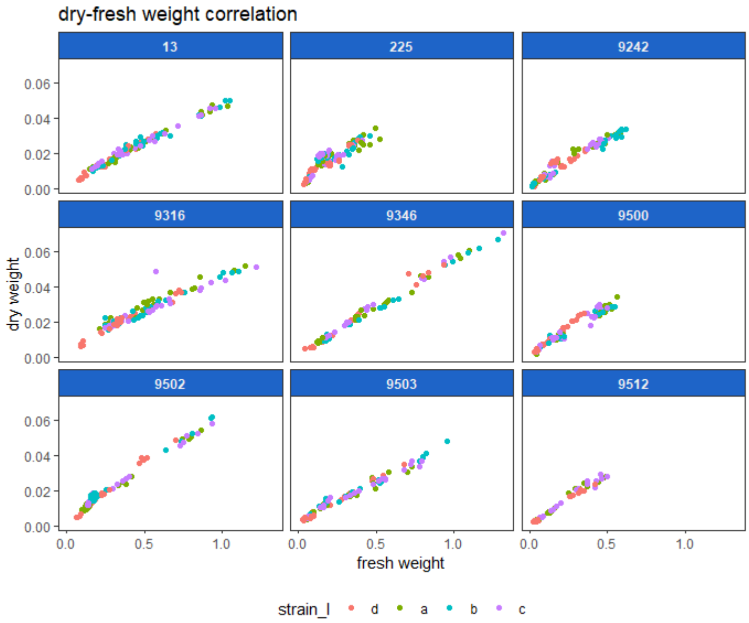
dry-fresh weight correlation
13
225
9242
9316
9346
9500
9502
9503
9512
dry weight
fresh weight
strain_l
d
a
b
c

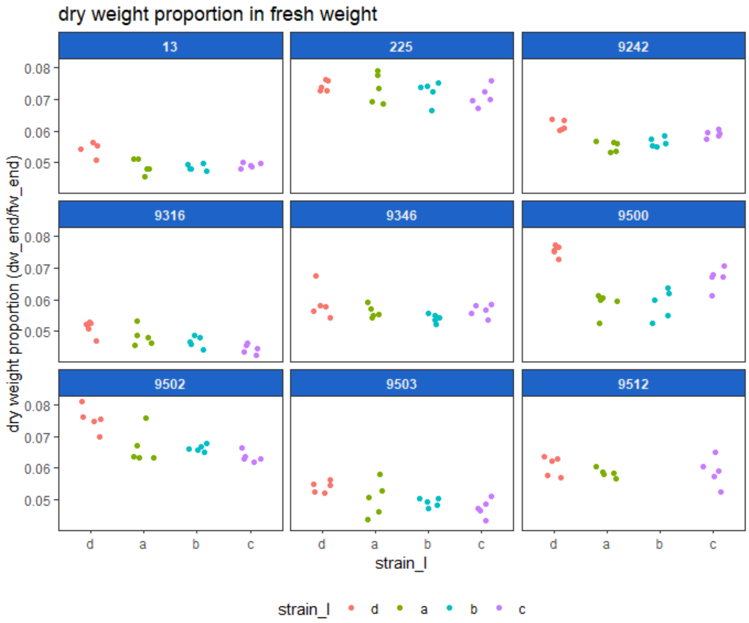
dry weight proportion in fresh weight
13
225
9242
9316
9346
9500
9502
9503
9512
dry weight proportion (dw_end/fw_end)
strain_l
d
a
b
c

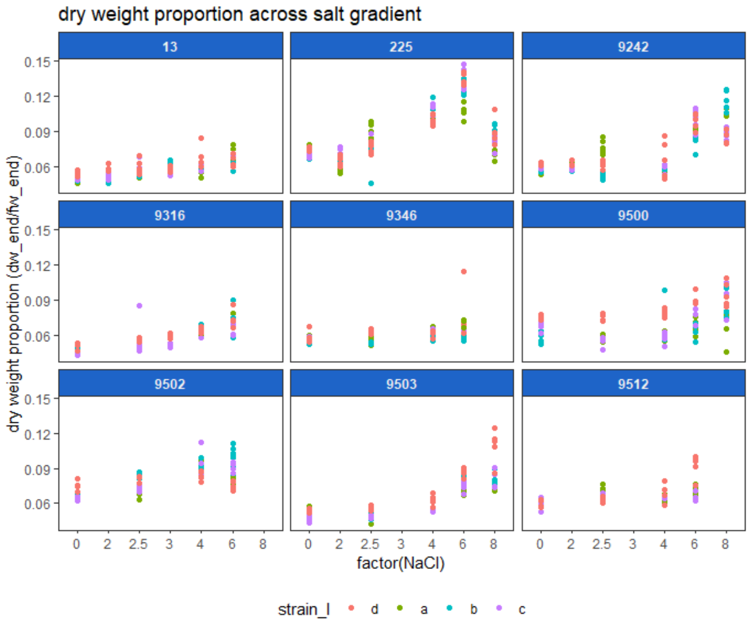


*Figure S13: (1) relation of dry weight to fresh weight, as measured at the end of the growth test period, (2) dry weight proportion (of wet weight) of three independent colchitetraploid strains (a, b, c) compared to their progenitor diploid strain (d, in red) for nine different genotypic backgrounds, and (3) dry weight proportion across the salt gradient.*
